# Scaling the Discrete-time Wright Fisher model to biobank-scale datasets

**DOI:** 10.1101/2023.05.19.541517

**Authors:** Jeffrey P. Spence, Tony Zeng, Hakhamanesh Mostafavi, Jonathan K. Pritchard

**Affiliations:** Department of Genetics, Stanford University; Department of Biology, Stanford University

## Abstract

The Discrete-Time Wright Fisher (DTWF) model and its large population diffusion limit are central to population genetics. These models describe the forward-in-time evolution of the frequency of an allele in a population and can include the fundamental forces of genetic drift, mutation, and selection. Computing likelihoods under the diffusion process is feasible, but the diffusion approximation breaks down for large sample sizes or in the presence of strong selection. Unfortunately, existing methods for computing likelihoods under the DTWF model do not scale to current exome sequencing sample sizes in the hundreds of thousands. Here we present an algorithm that approximates the DTWF model with provably bounded error and runs in time linear in the size of the population. Our approach relies on two key observations about Binomial distributions. The first is that Binomial distributions are approximately sparse. The second is that Binomial distributions with similar success probabilities are extremely close as distributions, allowing us to approximate the DTWF Markov transition matrix as a very low rank matrix. Together, these observations enable matrix-vector multiplication in linear (as opposed to the usual quadratic) time. We prove similar properties for Hypergeometric distributions, enabling fast computation of likelihoods for subsamples of the population. We show theoretically and in practice that this approximation is highly accurate and can scale to population sizes in the billions, paving the way for rigorous biobank-scale population genetic inference. Finally, we use our results to estimate how increasing sample sizes will improve the estimation of selection coefficients acting on loss-of-function variants. We find that increasing sample sizes beyond existing large exome sequencing cohorts will provide essentially no additional information except for genes with the most extreme fitness effects.

## 1 Introduction

The Discrete-Time Wright Fisher (DTWF) model and its large population limit the Wright-Fisher diffusion (WF diffusion) are workhorses of population genetics [1, 2]. These forward-in-time models describe the evolution of the frequency of an allele in a population, and can incorporate mutation, selection, and genetic drift.

Beyond providing a useful conceptual framework, the DTWF model and the WF diffusion enable inference of evolutionary parameters from data. A notable example is the Poisson Random Field (PRF) model [3] which relates the distribution of allele frequencies at a single site to the probability of observing a given number of sites where an allele is at a particular frequency in the sample (the site frequency spectrum; SFS). The SFS can be estimated from sequencing data, and hence the PRF provides a probabilistic model relating evolutionary parameters to observable genetic data. With a probabilistic model in hand, we can infer these evolutionary parameters by using standard techniques from statistical inference, such as maximum likelihood. This approach has been used throughout population genetics to infer population sizes [4], complex demographic models [5], and distributions of selection coefficients [6].

Unfortunately it is difficult to compute the distribution of frequencies at a single site for non-equilibrium demographies or models with natural selection under both the DTWF model and the WF diffusion. This distribution (potentially conditioned on being segregating at present) is one of the key ingredients of the PRF model. To illustrate these difficulties we will focus on the case of a single site with two potential alleles, *A* and *a*, in a panmictic monoecious haploid population.

Here our overarching goal will be to compute the likelihood of observing a particular allele frequency at present given various evolutionary parameters such as past population sizes, mutation rates, and selection coefficients. Throughout, we will use the notation **v**_*t*_ to represent a vector of these likelihoods at generation *t*. That is, entry *i* of **v**_*t*_ is the likelihood of observing exactly *i* copies of the *A* allele in the population at generation *t*. Thus, if we say that the present is generation *T*, our goal is to compute **v**_*T*_.

The simplest approach to computing these likelihoods is a naive forward-in-time application of the DTWF transition matrix. In this approach one assumes that at some point in the past the population is at equilibrium. Let **M**_eq_ ∈ ℝ^(*N* +1)*×*(*N* +1)^ be the Markov transition matrix describing the DTWF process for these ancient times. That is, for a population of size *N*, entry 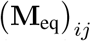 is the probability of transitioning from *i* individuals having the *A* allele in one generation to *j* individuals having the *A* allele in the subsequent generation. Note that **M**_eq_ depends on all of the evolutionary parameters of interest, such as population size, mutation rates, and selection. To obtain the equilibrium allele frequency distribution one can numerically solve the matrix equation

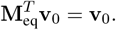

Thus, **v**_0_ is our vector of likelihoods at some point in the past, that we call generation 0. Once this equilibrium solution has been found, it can be integrated forward with changing population sizes or changing selective strengths by repeatedly applying the corresponding DTWF transition matrix for that generation. That is, supposing that the population is at equilibrium at generation 0, then to obtain the distribution at generation 1, one takes the transition matrix for that generation, **M**_0_, and then computes:

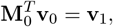

where **v**_1_ is now a vector containing the probabilities of observing each of the different possible allele frequencies at generation 1. This may be repeated until we arrive at the present day, allowing us to compute the probability of observing a given number of *A* alleles in the population at present. If we do not sample the whole population, then we can obtain the probability of a given sample frequency by sampling from the population without replacement.

The crux of this naive method is repeated matrix-vector products; we repeatedly multiply the vector of probabilities for a given generation by the transition matrix for that generation. Matrix-vector products for an (*N* + 1) ×(*N* + 1) matrix take *O*(*N* ^2^) time, and so this approach requires *O*(*N* ^2^) time to compute likelihoods if one has access to the stationary distribution. Given that for humans the present day effective population size may be in the millions (or more) [7], this naive approach is obviously not scalable.

To avoid the onerous *O*(*N* ^2^) runtime, many approaches are based on the idea that population sizes are usually quite large, and as such one might consider a large population size limit of the DTWF model. This limit assumes a fixed sample size *n* and then takes *N* to infinity, although in practice this approach is used for all but the smallest population sizes. The resulting continuum limit is the celebrated WF diffusion [2]. This continuum limit is still a Markov process, but instead of having a finite state space, the state space is the continuous unit interval [0, 1] of allele frequencies. As such, the limiting process is no longer a discrete time, discrete space Markov chain, but a continuous time, continuous space Markov chain whose evolution is described by a stochastic differential equation. This stochastic differential equation then allows one to write a partial differential equation (PDE) that describes how the likelihood of observing different allele frequencies changes over time. Similar to the naive DTWF approach, one may solve this PDE at equilibrium at some time in the ancient past and then evolve the likelihoods forward in time to obtain the likelihoods at the present. The advantage of this approach is that whether the population has ten thousand individuals or ten million individuals the PDE is functionally the same. As a result, the runtime of computing likelihoods becomes independent of *N*. This approach has been extremely fruitful, resulting in numerical solutions [5] and spectral approaches [8, 9, 10]. Yet, numerically solving a PDE can be difficult and error-prone, and some methods have been found to return negative “probabilities” [11]. On the other hand, spectral methods can approximate the transition density from one allele frequency to another very accurately, but require computationally costly symbolic algebra.

Another line of work takes a backward-in-time approach using ideas from Kingman’s Coalescent [12], which results in likelihoods equivalent to those computed forward-in-time using the WF diffusion [13] (but see [4, 14] for coalescent approaches that are equivalent to the DTWF model). These backward-in-time approaches have the advantage of only scaling in terms of the sample size, *n*, as opposed to the population size, *N*. Usually, *n* ≪ *N*, and so this scaling can result in substantial computational speedups. For example, Polanski and Kimmel developed an approach to compute likelihoods under the coalescent for arbitrary past population size functions in *O*(*n*^2^) time [15]. A major downside of coalescent approaches is the difficulty of incorporating natural selection [16]. One of the keys of coalescent approaches is that the genealogical process can be separated from the mutational process — one can first sample a genealogy and then drop mutations along the branches of the sampled tree to obtain the alleles of the present day individuals. This separation of genealogy and mutation implicitly rests on the assumption that an individual’s fitness is independent of their allelic type, which is violated when there is natural selection. One can in principle obtain a genealogical process in the presence of natural selection [16], but inference under this process is generally intractable.

Forward-in-time and backward-in-time methods each have advantages and disadvantages, and so a number of hybrid approaches have been developed. For example, `momi` [11, 17] uses the fact that the genealogies of the backward-in-time coalescent can be embedded in a forward-in-time Moran model. `momi` can model complex demographies, but cannot model selection. A similar trick is used in `moments` [18], but `moments` can model natural selection while remaining a good approximation to the WF diffusion. oh While the WF diffusion and coalescent can enable more efficient inference, they are only accurate for sufficiently common alleles. This inaccuracy has been noted several times, usually in the context of the coalescent, but the coalescent and WF diffusion are dual processes, so these inaccuracies also apply to the WF diffusion. There begin to be notable discrepancies between the DTWF model and the WF diffusion for the likelihood of observing rare alleles in the sample when the sample size, *n*, is larger than roughly the square root of the population size, 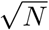 [14, 19, 20, 21, 22].

The diffusion limit also assumes that all relevant evolutionary parameters such as mutation rates and strengths of selection scale like 1*/N* in the limit [2]. That is, if selection is ≫1*/N*, then the diffusion approximation breaks down [20].

Most of the methods discussed above compute likelihoods under processes equivalent to the WF diffusion, potentially suffering from these problems. Bhaskar, Clark, and Song introduced a coalescent approach dual to the DTWF model that scales like *O*(*n*^3^) but cannot incorporate natural selection [19]. More recently, Krukov and Gravel developed an approach that can model natural selection using additional bookkeeping to accurately compute likelihoods under the DTWF process [20]. Unfortunately, this approach scales like *O*(*n*^4^).The distinction between the DTWF process and the WF diffusion becomes apparent when *n* is larger than 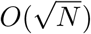. In this regime, a runtime of *O*(*n*^4^) implies a runtime of at least *O*(*N* ^2^), no better than the naive forward-in-time approach using the DTWF transition matrix.

There has been much interest in determining the extent to which natural selection acts against loss-of-function variants in each gene in the human genome using massive exome sequencing datasets [23, 24, 25, 26, 27, 28]. This regime — sample sizes in the hundreds of thousands, and extremely strong selection — is exactly where differences between the DTWF model and the diffusion become most pronounced, highlighting the need for more computationally efficient methods.

Here we reconsider the naive approach of using the forward-in-time DTWF process. While the most basic method of computing likelihoods using the DTWF process costs *O*(*N* ^2^) time, we show that transition matrices under a broad class of DTWF processes can be replaced by highly structured matrices enabling likelihood computations in *O*(*N*) time, while having a provably small approximation error. We obtain a similar speedup for Hypergeometric sampling that may be of independent interest. We provide a high level overview of our approach in Section 2, and in Section 3.1 we show empirically that our approach is highly accurate and can scale to sample sizes in the billions. In Section 3.2 we use our approach to explore the utility of using loss-of-function variants to estimate selection coefficients in large samples. We find that increasing sample sizes beyond current values will provide little value for estimating the selection coefficients of most genes, and will only prove useful for estimating extremely strong selection coefficients. We discuss the limitations and future directions for our approach in Section 4. Formal proofs are deferred to Appendix A. We apply our approach in an empirical Bayes framework to estimate the strength of selection against loss-of-function variants using large-scale exome sequencing data in a companion paper [29]. Software with a python API implementing our approach is available at https://github.com/jeffspence/fastDTWF.

## 2 Overview of approach

Throughout we focus on a single locus with two alleles, *A* and *a*. Our goal is to compute the likelihood of observing a given frequency of the *A* allele at a particular locus in a sample from a population evolving according to the DTWF model. All of the evolutionary forces present in a general DTWF model are captured by the transition matrices, **M**_*t*_. In particular, **M**_*t*_ is affected by the population sizes in generations *t* and *t* + 1, as well as the mutation rates and effects of selection. For general non-equilibrium populations where these evolutionary parameters are changing over time, a naive approach to computing these likelihoods involves three steps:

1. We assume that at some point in the past the population was at equilibrium, and we compute **v**_0_, a vector with *N* + 1 entries, indexed from 0 to *N*, where entry *i* is the probability of observing *i* copies of the *A* allele in the population at equilibrium, with *N* being the population size.
2. We evolve these probabilities forward according to the DTWF transition matrix for each generation, until we reach the present. That is, for generation *t* − 1, let **M**_*t*−1_ be the DTWF transition matrix. Then, (**M**_*t*−1_)_*i,j*_ is the probability of going from *i* copies of the *A* allele in the population in generation *t*− 1*s* to *j* copies of the *A* allele in generation *t*. To obtain the probability of each allele frequency in the population at generation *t* we can compute

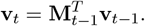 Say that we want to compute the likelihood at generation *T*. Then, given **v**_0_, we can compute the population-level allele frequency probabilities as

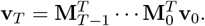
3. The first two steps compute the probability of observing each different possible allele frequency in the *population*, and so we still must obtain the probability of observing each different possible allele frequency in a *sample* from the population. Let **S** be a matrix where entry **S**_*i,j*_ is the probability of seeing *j A* alleles in a sample given that there are *i A* alleles in the population. We may therefore obtain the probabilities of observing each possible allele frequency in the sample, **v**_sample_ as

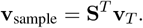

Each of these steps is intractable using a naive approach because they rely on matrix-vector multiplication, requiring *O*(*N*^2^) time. Yet, if we were able to make matrix-vector multiplication much faster, then this approach suddenly becomes attractive — it is conceptually straightforward; easy to extend to incorporate selection and changes in mutation rates or population sizes; and numerically stable because all of the entries in all of the vectors and matrices are positive, avoiding the catastrophic cancellation that plagues some other approaches.

Our approach is to replace the DTWF transition matrices **M**_*t*_ and the sampling matrix **S** with approximate versions, 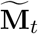 and 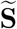 that allow for matrix-vector multiplication in *O*(*N*) time, while guaranteeing that 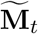 and 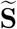 are extremely “close” to **M**_*t*_ and **S** respectively.

Our main results are about matrices where each row consists of the *N* + 1 entries of a probability mass function of a Binomial distribution with sample size *N*. We call this class of matrices Binomial transition matrices. While this class of matrices may seem esoteric, the transition matrices of many types of DTWF models are either themselves Binomial transition matrices or can be well-approximated using Binomial transition matrices.

On an intuitive level, our results rely on the observation that all of the “action” in a Binomial distribution happens on the scale of 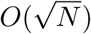. This observation manifests itself in two distinct ways, each allowing us to speed up matrix-vector multiplication by a factor of 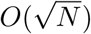. The first is that with overwhelming probability a Binomial random variable will fall within 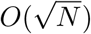 of its mean. In our setting this allows us to ignore all of the entries in each row of the transition matrix, except for 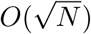 entries around the mean, while only introducing a small amount of error.

The second way in which the “action” of Binomial distributions happens at the 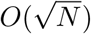 scale is somewhat more subtle. If two Binomial distributions with the same sample size have success probabilities that are very close to each other, then the resulting distributions are very similar. More precisely, if one has a binomial distribution with sample size *N* and success probability *p*, and wants to find another Binomial distribution with sample size *N* that is no further than *ε* distance away (measured in total variation distance) then one can choose any Binomial distribution with a success probability up to 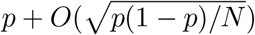. Ultimately, this means that many rows of a Binomial transition matrix must be very smilar. As a result, we may replace the matrix by one with only 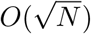 unique rows, while only incurring a small amount of error per row.

These two tricks together result in a matrix that is both highly structured — it is extremely low rank with only 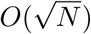 unique rows — and highly sparse — each row only has 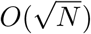 non-zero entries. Below, we will provide more intuition about these two tricks, and also describe how together they allow for computing matrix-vector products in *O*(*N*) time.

To obtain sampling probabilities, we still need to be able to perform a matrix-vector product with the sampling matrix **S**. If we obtain our sample from the population by sampling without replacement as is usually assumed, then if we know that there are *K* copies of the *A* allele in the population of *N* individuals, the number of *A* alleles in a sample of size *n* is Hypergeometric distributed with parameters *N, n* and *K*. That is, just as the rows of **M** are the probability mass functions of Binomial distributions, the rows of **S** are the probability mass functions of Hypergeometric distributions. Surprisingly, the same properties of Binomial distributions that allowed us to perform fast matrix-vector products are also true of Hypergeometric distributions. This allows us to use very similar tricks to quickly compute matrix-vector products with the sampling matrix, **S**.

In contrast, if one obtains a sample via sampling *with* replacement, then that sampling process can be represented as one additional generation of a DTWF process, but with no mutation and no selection. One can also model sampling with replacement where the sampling is biased toward individuals with one allele or the other, in which case the sampling process is equivalent to a single generation of the DTWF model without mutation, but with a particular form of natural selection. More complicated sampling processes (e.g., sampling without replacement, but biased toward one allele or another) may be possible to treat using our techniques, but would require additional considerations beyond those presented here.

### 2.1 Total variation distance and matrix norms

Our results about Binomial and Hypergeometric distributions are in terms of total variation distance, an important metric on the space of distributions. For the discrete distributions taking values in 0, 1, 2, …, *N* that we consider here, total variation distance is simply half the 𝓁_1_ distance between the probability mass functions. That is, for a distribution *P* and a distribution *Q*, we write the total variation distance between them, *d*_*T V*_ (*P, Q*), as

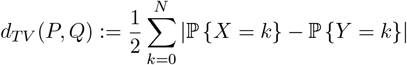

where *X* is a *P* -distributed random variable and *Y* is a *Q*-distributed random variable.

We present our results in terms of total variation distance because it is very closely related to a particular matrix norm. The *1-operator norm* of a matrix, which we denote by ‖ · ‖_1_, is defined as

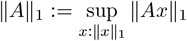

for any matrix *A*, where ‖ · ‖_1_ applied to vectors is the usual 𝓁_1_ norm (i.e., the sum of the absolute values of the entries). Note that the 1-operator norm is *not* the sum of the absolute values of the entries of the matrix. In fact, the 1-operator norm is the maximum of the column-wise 𝓁_1_ norms. The proof of this well-known result is included in Appendix D for completeness.

The reason we are interested in the 1-operator norm is because it allows us to bound how much error we might introduce by replacing a DTWF transition matrix by an approximation. In particular, if we have a DTWF transition matrix **M** and we can construct an approximate matrix 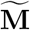 such that 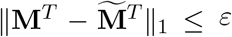, then when we compute the matrix-vector products required for computing likelihoods, we will have that 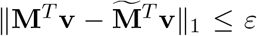. Since the rows of a DTWF matrix correspond to probability mass functions, we see that bounding an approximation’s row-wise 𝓁_1_ distance is equivalent to bounding the total variation distance between the corresponding distributions up to a factor of 2.

### 2.2 Binomial transition matrices are approximately sparse

The first key for our approach is straightforward: Binomial random variables are very unlikely to be too far away from their means. A Binomial random variable will be more than

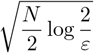

away from its mean with probability less than *ε*. This is a celebrated result due to Hoeffding [30], and shows that with overwhelming probability, a Binomial random variable will be within a constant factor times 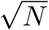 of its mean. In turn, this implies that we can ignore all but 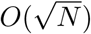 entries in each row of a Binomial transition matrix while only incurring a constant (in *N*) total variation distance. This property of Binomial distributions is illustrated in Figure 1A.

**Figure 1:**
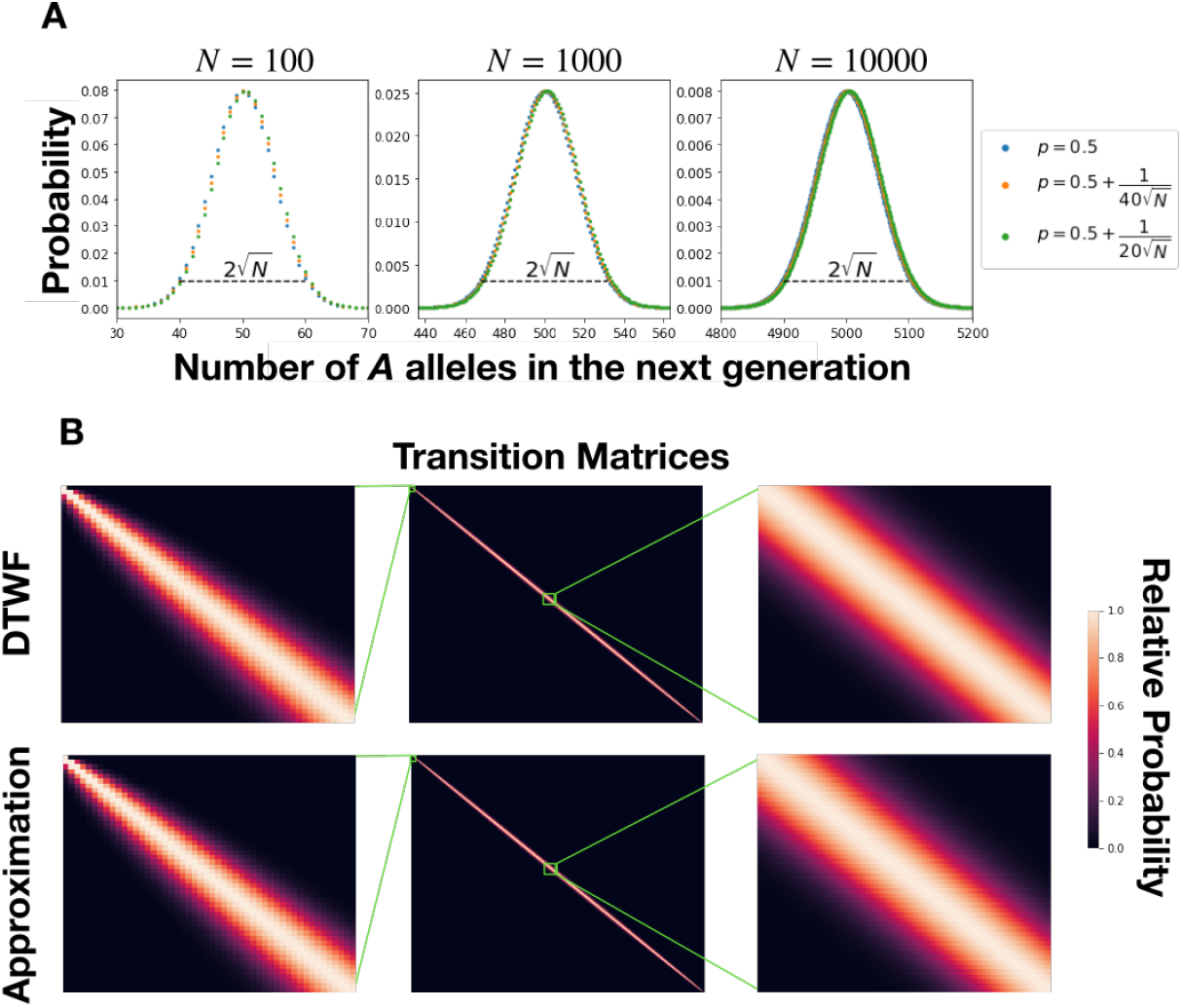
**A** Probability mass functions for Binomial distributions across a range of values of *N*. Most of the mass is contained within 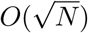 of the mean, and distributions with success probabilities *p* within a small factor of 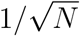 of each other are virtually indistinguishable. **B** The transition matrix of the neutral DTWF process with *N* = 5000 as well as our approximation of that matrix represented as a heatmap. Rows are normalized so that the maximum of each row is 1, and regions from the top left and middle are expanded. The results are nearly indistinguishable, except that there is very subtle horizontal banding near the middle of the transition matrix resulting from having nearby rows be copies of each other.

This simple observation alone provides substantial savings in terms of memory and runtime for computing matrix-vector products, which has been noted previously [31, 32], as we can simply take each row of a Binomial transition matrix, and replace all of the entries that are too far away from the corresponding mean of the row by zero. We can choose the point at which we begin setting entries to zero to obtain a given error tolerance, *ε*. To ensure that the resulting approximate matrix is still a valid stochastic matrix (i.e., the rows still sum to one and hence are valid probability distributions) we divide the remaining non-zero entries by their sum, which by construction perturbs them by a multiplicative factor no larger than 1*/*(1 − *ε*).

As a concrete example, to capture all but 10^−16^ of the probability in a Binom<u>i</u>al distribution, corresponding roughly to the limits of numerical precision, we can ignore all but 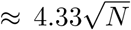 entries on either side of the mean when computing matrix-vector products. This means that we can approximate the matrix as having only 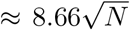 non-zero entries in each row, resulting in a theoretical speedup by a factor of 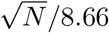. When *N* is 1000, this corresponds to a factor of ≈7.3× speedup over the naive approach. When *N* is 10 million, the speed up is ≈730×. This high degree of sparsity is visually apparent in the 5000× 5000 neutral DTWF transition matrix shown in Figure 1B.

### 2.3 Binomial transition matrices are approximately low rank

The second key for our approach is more subtle, and relies on the fact that Binomial distributions with similar success probabilities have similar distributions in terms of total variation distance. This makes sense on an intuitive level — flipping *N* coins that come up heads with probability *p* should result in a similar distribution of outcomes to flipping *N* coins that come up heads with probability *p* + δ. What is less obvious is the length scale at which this occurs. That is, how large can δ be in terms of *p* and *N* while still keeping the total variation distance between the two distributions below a specified level? The answer is 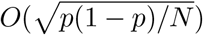, which is visually apparent in the different colored points in Figure 1A, and which we prove in Appendix A. In practice, we use 0.1 as an empirical cutoff for determining what distributions are similar — that is, we consider distributions with success probabilities *p* and 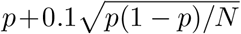 to be close enough to be treated as identical. We now consider partitioning the unit interval [0, 1] into blocks such that for *p* and *p*′ in the same block, two Binomial distributions with size *N* and success probabilities *p* and *p*′ will have total variation distance less than *ε*. We show in Appendix A that we can achieve such a partitioning of [0, 1] with 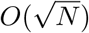 blocks.

We can use this partitioning to approximate a Binomial transition matrix by an extremely low rank matrix, while bounding the 𝓁_1_ error introduced to each row. For each separate block in the partition of [0, 1], we can pick a representative success probability. Since there are 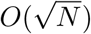 blocks in this partition, we end up with 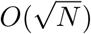 representative success probabilities. Then, for each row of the Binomial transition matrix, we can consider its success probability, determine which block of the partition it is in, and replace that row by the probability mass function of the Binomial random variable with the corresponding representative success probability. Because the original row, and the new row correspond to probability mass functions for Binomial distributions that are close in total variation distance, the rows are close in 𝓁_1_ distance. After replacing each row, the resulting matrix can have at most 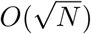 unique rows, meaning that for large *N* it is extremely low rank. Additionally, since each row of the resulting matrix is still the probability mass function of some Binomial distribution, the resulting matrix is still a Binomial transition matrix.

While picking an arbitrary success probability for each block in the partition of [0, 1] bounds the total variation distance, different choices can results in different accuracies of the approximation over repeated matrix-vector multiplications. For instance, if one were to choose the smallest success probability within each block, then, for a neutral DTWF transition matrix, the expected frequency in the next generation would never be larger than the current frequency and would often be slightly smaller. Over evolutionary time-scales this would act similarly to negative selection, affecting the long-term accuracy. Instead of choosing arbitrarily, we found that in practice a moment-matching approach is extremely accurate. Briefly, when performing matrix-vector multiplication with a non-negative vector **v**, for a given block of the partition of [0, 1] we use the weighted average of the success probabilities in **M** that fall within that block with weights proportional to the corresponding entries of **v**.

Specifically, suppose that rows *i, i* + 1, …, *j*, with success probabilities *p*_*i*_, *p*_*i*+1_, …, *p*_*j*_ will all be represented by a single row with success probability 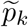. If we assume that the success probabilities are ordered, then setting 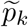 to be any value between *p*_*i*_ and *p*_*j*_ will bound the total variation distance, but some choices may result in worse long-term accuracy. If we are approximating **M**^*T*^ **v** for some **v** with nonnegative entries, then we set

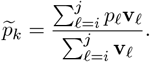

This choice guarantees that the expected frequency in the next generation of an allele with frequency in the current generation chosen with probabilities proportional to **v** is matched between the true and approximate processes. In the event that the denominator is exactly 0, then the choice of 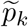 does not matter as we will see in the next section.

### 2.4 An *O*(*N*) algorithm for approximately computing matrix-vector products for Binomial transition matrices

These two ingredients — that each row of a Binomial transition matrix is close in 𝓁_1_ distance to a row with only 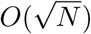 non-zero entries, and that each row of a Binomial transition matrix is close in 𝓁_1_ distance to the corresponding row of a Binomial transition matrix with only 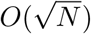 unique rows — are sufficient to derive a substantially faster algorithm for approximately computing (transposed) matrix-vector products.

The key idea is to replace the original Binomial transition matrix **M** by an approximation 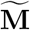, which we construct by first choosing a Binomial transition matrix with 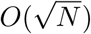 unique rows that is close in row-wise 𝓁_1_ distance to **M** and then sparsifying each row of that matrix. If we perform each of these steps so that they introduce a row-wise 𝓁_1_ distance of at most *ε/*2, the triangle inequality implies that the two steps together introduce a total row-wise 𝓁_1_ error of at most *ε* per row.

Once we have our approximate matrix, we can quickly perform matrix-vector multiplications (Figure 2). The algorithm involves noting that computing 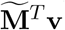 can be thought of as first multiplying each row of 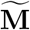 by the corresponding entry of **v**, and then summing up those resulting vectors. That is, one first performs *N* + 1 scalar-vector multiplications, and then sums up the *N* + 1 resulting vectors. Our speedups come from two places.

**Figure 2:**
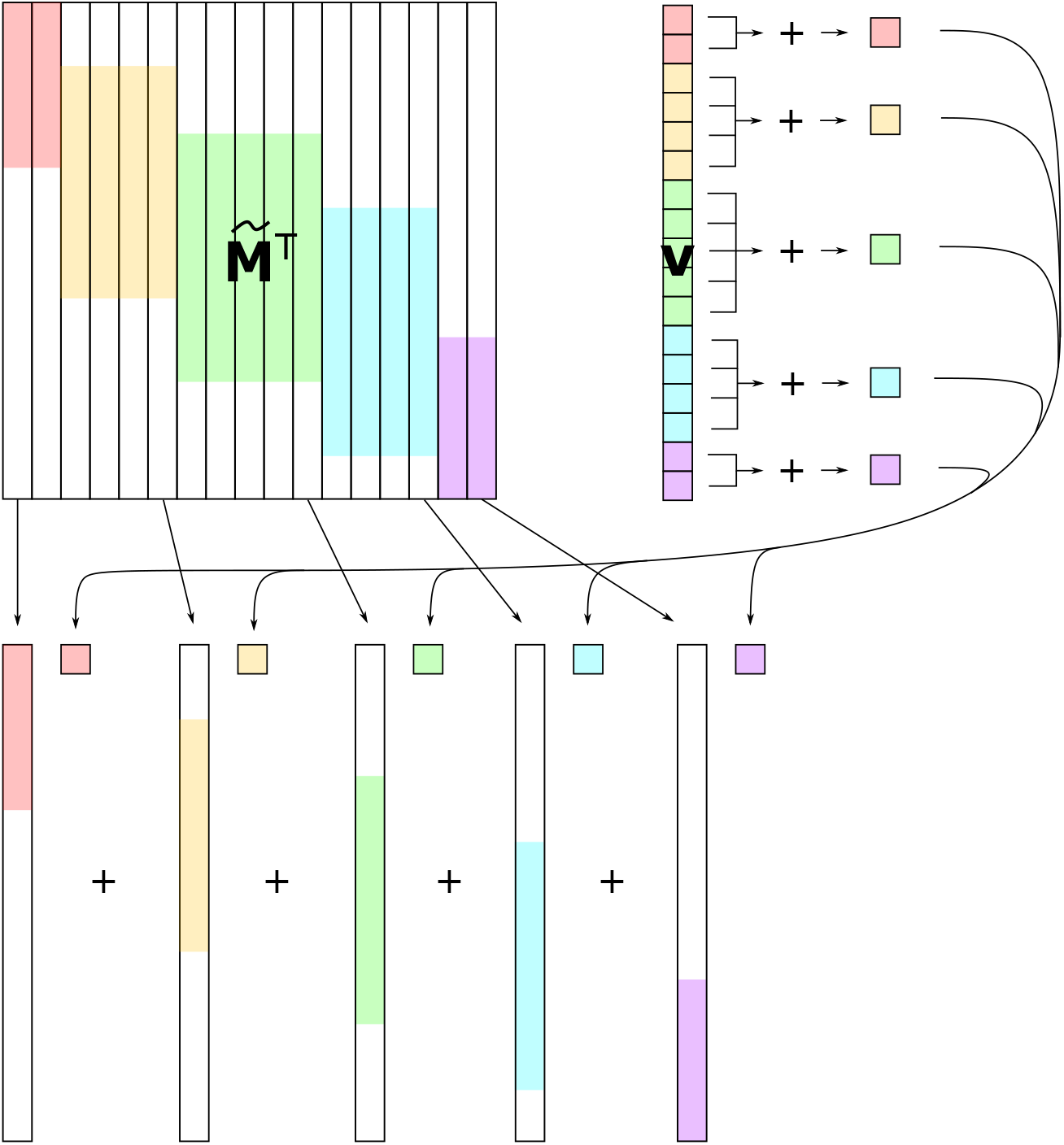
Schematic of fast matrix-vector multiplication algorithm. White regions of 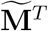 correspond to zeros, colors correspond to columns of 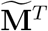 (i.e., rows of 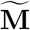) that are identical, and the corresponding entries of **v**. The algorithm proceeds by first combining entries of **v** that correspond to identical rows of **m**, then multiplying the resulting scalars by the representative rows of 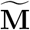, using sparse scalar-vector multiplication. The resulting sparse vectors are then summed using sparse vector addition.

First, since 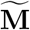 has 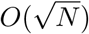 unique rows, many rows must be identical. Instead of multiplying each identical row by the corresponding entry of **v** and then summing, we can instead first sum up all of the entries of **v** that correspond to identical rows, and then multiply one representative row by this sum of the relevant entries of **v**. This observation means that after grouping the entries of **v**, we only need to perform 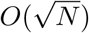 scalar-vector multiplications, and then sum up the 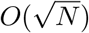 resulting vectors.

Second, since each row of 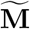 is sparse, we can ignore all of the zero entries when performing scalar-vector multiplication and vector addition. Our vectors only have 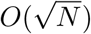 nonzero entries, making both of those operations cost 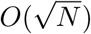 time. Overall, this means that we must perform 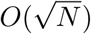 operations, each taking 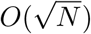 time, resulting in a runtime of *O*(*N*). We give a visual depiction of our algorithm in Figure 2. Details and technical proofs are presented in Appendix A.

There are a few technical details and assumptions in achieving a truly *O*(*N*) runtime. First, we obviously cannot store or even look at each entry in **M**, because doing so would require *O*(*N* ^2^) space and time. Instead we represent a Binomial transition matrix (or DTWF matrix) as simply the *N* + 1 success probabilities corresponding to each row. We can then represent 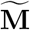**s** by storing only the locations and values of the 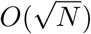 entries for each of the 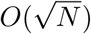 unique rows. Second, determining which representative row to use for each row of **M** can — in the worst case — require *O*(*N* log *N*) time; one can keep the break points of the partition of [0, 1] in an ordered list, and then for each row of **M** one must take its corresponding success probability and search through the sorted list of break points. This binary search requires *O*(log *N*) time for each row, resulting in a runtime of *O*(*N* log *N*). Instead, we assume that the rows of **M** are ordered in terms of success probabilities. In Appendix A we present a simple *O*(*N*) algorithm for assigning ordered success probabilities to blocks of the partition. In the DTWF context, this ordering corresponds to the case where the expected allele frequency in the next generation is non-decreasing in the current allele frequency. This assumption is biologically reasonable, and would only be violated by something like extreme and unusual density-dependent selection. Indeed, this assumption holds for standard models of haploid or diploid selection and mutation.

### 2.5 Efficiently obtaining the likelihood of a sample from population likelihoods

Using the algorithm in the previous section allows us to compute population-level likelihoods. In general, we do not have access to population-level data and must instead obtain a sample from the population, which we assume is done uniformly at random without replacement (i.e., simple random sampling). If we take a sample of size *n*, then supposing that there are *K* copies of the *A* allele in the population, the number of *A* alleles in the sample is Hypergeometric distributed with parameters *N, n*, and *K*. Thus, to obtain the probability of observing a given number of *A* alleles in the sample, we must take an average of Hypergeometric distributions weighted by the probability of having a given number of *A* alleles in the population. We can write this as a matrix equation, using an (*N* + 1)×(*n* + 1) dimensional sampling matrix **S** whose *K*^th^ row is the probability mass function of a Hypergeometric random variable with parameters *N, n*, and *K* (assuming that the rows are 0-indexed). If the probabilities of observing 0, 1, …, *N* copies of the *A* allele are contained in the *N* + 1 dimensional vector **v**, then we can obtain the vector, **v**_sample_ of probabilities of observing 0, 1, …, *n* copies of the *A* allele as

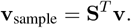

Naively computing this matrix-vector product would take *O*(*nN*) time and space, which is prohibitive for large sample sizes.

Somewhat surprisingly, if we assume that the sample size is not too large as a function of the population size — i.e., *n* ≤ *αN* for any fixed *α* < 1 as *N* grows, then the matrix **S** has properties very similar to a Binomial transition matrix despite having Hypergeometric rows instead of Binomial rows. On the one hand, the difference between the Hypergeometric distribution and the Binomial distribution is the same as sampling with and without replacement. Thus, for samples that are small, we might expect that it would be rare to sample the same individual twice when sampling with replacement, and so sampling with and without replacement should be similar. On the other hand, our results apply even as the sample size grows with the population size, and even for cases where, for example, we are sampling 99% of the population. In such cases, when sampling the same number of individuals, but *with* replacement we would almost certainly resample some individuals multiple times, and so it is surprising that sampling with and without replacement might have similar properties in this regime. Yet, as we show in Appendix A, **S**^*T*^ is close in 1-op<u>e</u>rator norm to a highly structured matrix 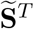 such that 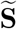 has 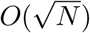 unique rows and each row of 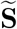 has 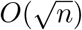 non-zero entries. The proof of this result relies on similar considerations as the Binomial case but applied to Hypergeometric distributions. Together, these two properties were all that was required to obtain our *O*(*N*) algorithm for (transposed) matrix-vector products, and so we may use exactly the same trick to compute **S**^*T*^ **v**.

As above, some care needs to be taken when choosing a row as a “representative” of a set of similar rows. In this case, we again use a moment matching approach, taking a mixture of two Hypergeometric distributions so that the expected value of an allele chosen uniformly at random with probabilities proportional to **v** is matched between the approximate and real sampling matrices.

This completes our overview of our approach to approximately computing likelihoods. By applying our algorithm to the DTWF transition matrix, we can efficiently compute the stationary distribution, and integrate that distribution forward to the present, obtaining the present-day population-level likelihood. Then, by applying essentially the same algorithm to the sampling matrix, we can efficiently obtain sample-level likelihoods.

## 3 Numerical Results

In this section we present numerical results about the runtime and accuracy of our method as well as an application to how selection and demography interact to affect the distribution of observed frequencies in large sample sizes. These results have implications for how much information we might hope to gain about selection coefficients as sample sizes grow.

Before discussing the numerical results, we discuss some practical implementation details. The theoretical accuracy guarantees of our approach involve implicit constants. For example, we know that we can choose a 𝒸_*ϵ*_ such that if we replace a row of the transition matrix that corresponds to a Binomial with success probability *p*, with a row that corresponds to a success probability of 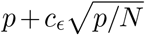, then we induce an 𝓁_1_ error that is bounded by *ϵ* regardless of *N* or *p*. Yet, our proof that such a constant exists is non-constructive (and our proof is such that making it constructive would result in a much smaller *c*_*ϵ*_ than necessary). As a result, we instead have the user specify two hyperparameters, which together determine both the runtime and accuracy. Regardless of their setting, our matrix-vector multiplication runs in *O*(*N*) time, but the hyperparameters determine the size of the constant hidden by the big-O notation, as well as the accuracy. The first hyperparameter is the *c*_*E*_ described above, which can alternatively be described as how many standard deviations away two rows’ success probabilities can be before we will not allow one to be a copy of the other. Setting this value to be smaller results in a longer runtime, but higher accuracy. Unless otherwise specified, we set this hyperparameter to be 0.1 for all analyses. The other hyperparameter determines how sparse to make the rows. Here our theoretical results *do* provide a constructive guarantee, so the user specifies a particular row-wise 𝓁_1_ error tolerance, and the rows are only sparsified to a level that guarantees a smaller error. Again, setting this parameter to be smaller results in longer runtimes, but higher accuracy. Unless otherwise specified, we set this sparsification error tolerance to be 10^−8^. We also use the same hyperparameters to specify the level of accuracy for the sampling matrix, **S**, but allow the user to set them to different values. Throughout, we always set the *c*_*ϵ*_ hyperparameter for the sampling matrix to 0.05 and the row sparsification hyperparameter to 10^−8^.

### 3.1 Runtime and accuracy

To begin, we confirm the linear runtime of our matrix-vector multiplication algorithm and compare our implementation to the runtime of the naive quadratic approach. For these analyses, we considered a neutral DTWF model with bidirectional recurrent mutation at a rate of 1.25×10^−8^ per generation. We compared the run-time of matrix-vector multiplication with this matrix and a random vector, where each entry is independent and identically distributed Uniform(0, 1), and then normalized to sum to one. Compared to the *O*(*N*^2^) naive matrix-vector multiplication algorithm, our approximate algorithm has a more favorable *O*(*N*) scaling (Figure 3**A**). While big-*O* notation hides constant factors, we see that across population sizes (from *N*≥1000) our approximate algorithm is faster than the naive algorithm. Indeed, at *N* = 1000, our approximate algorithm is slightly faster (6% speedup), while at *N* = 10,000,000, we predict that our algorithm would be about 215,000× faster, with the naive algorithm predicted to take over 75 *days* and our algorithm taking 31 *seconds* (we did not run the naive algorithm for *N* > 79,000 due to the prohibitive runtime). Similar results hold for matrix-vector multiplication using the sampling matrix (Figure 3**B**), where we obtain substantial speedups regardless of whether we sample 5% or 50% of the population.

**Figure 3:**
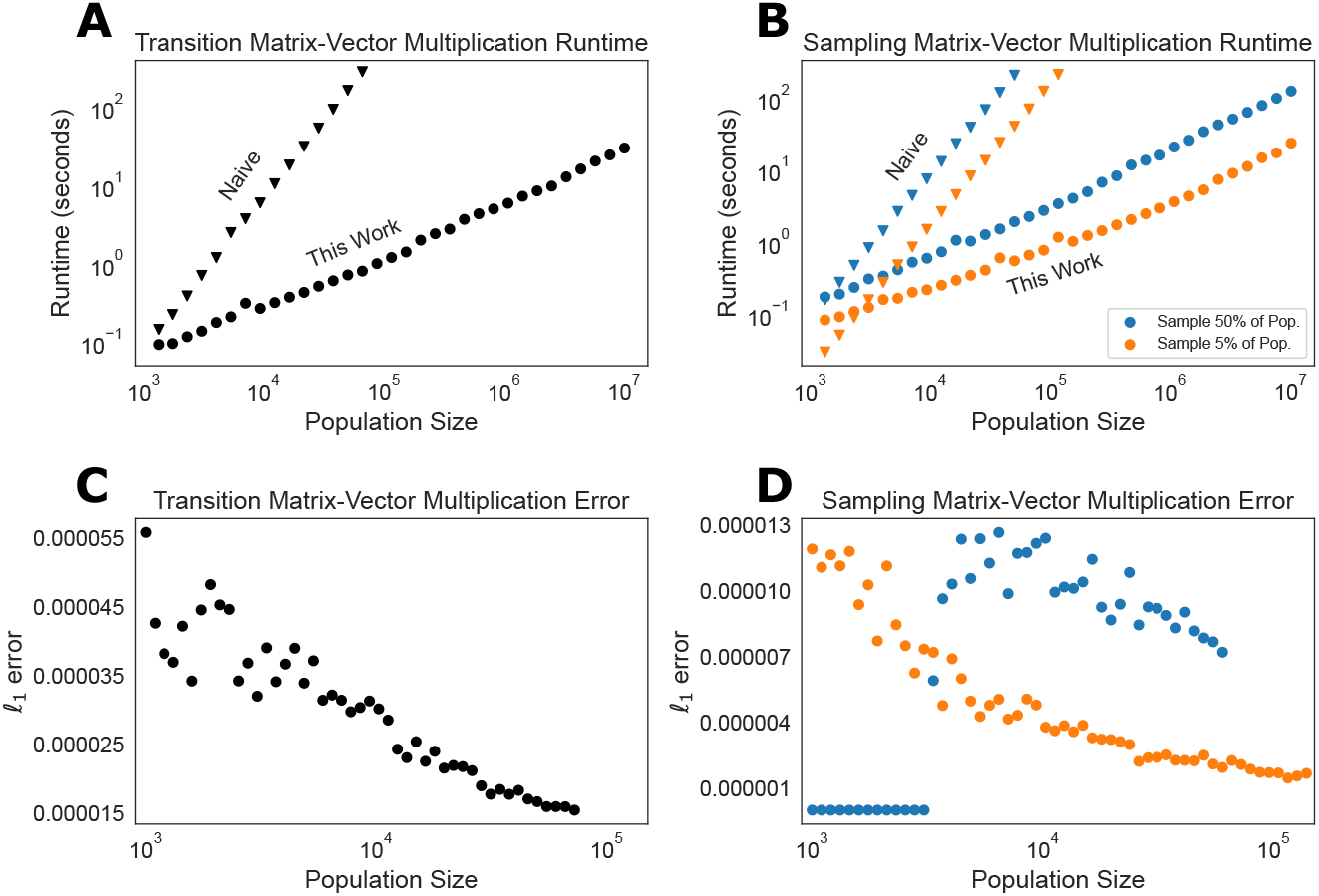
Runtime and accuracy of approximate algorithm. **A** Runtime of the approximate and naive matrix-vector multiplication algorithms for a DTWF transition matrix as a function of the population size, plotted on log-log scale. The naive algorithm scales quadratically, while the approximate algorithm proposed here scales linearly. **B** Runtime of the approximate and naive matrix-vector multiplication for the sampling matrix as a function of the population size, plotted on log-log scale. The runtime when the sample size is 50% of the population size is in blue, and orange is the runtime when the sample size is 5% of the population size. In both **A** and **B**, runs that were expected to take more than 5 minutes were not run. **C** The 𝓁_1_ error of the vector resulting from our approximate matrix-vector multiplication algorithm, compared to the vector obtained from exact matrix-vector multiplication for a DTWF transition matrix multiplied with a random vector. **D** Same as **C**, but for the sampling matrix, when considering sampling either 50% (blue) or 5% (orange) of the population. In both **C** and **D** it is apparent that the 𝓁_1_ error does not grow (and in fact decreases) with increasing population size, consistent with our theoretical guarantees.

Having confirmed that our approximate matrix-vector multiplication algorithm provides a substantial speedup, we turned to assessing its accuracy. We began by considering the accuracy of performing a single matrix-vector multiplication. That is, we considered the 𝓁_1_ error between the result of the exact matrix-vector multiplication algorithm, and the approximate matrix-vector multiplication algorithm, 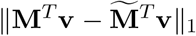. We used the same randomly generated **v** as we used for benchmarking the runtimes. Our theoretical results imply that even as *N* grows, our approximation should not get any worse. This theory is borne out empirically, where in fact we see that the error is small across all *N*, and the approximation actually appears to become more accurate as *N* gets large (Figure 3**C**).

We see similar results for the sampling matrix (Figure 3**D**), where the error is slightly larger when sampling a larger fraction of the population, but is low across all *N* and both of the sampling fractions considered. When the population size is small and we sample 50% of it, we see almost zero error. The reason for this is subtle, but by our construction of the approximate sampling matrix, 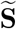 if no more than two rows are combined into any single representative row, then our algorithm exactly recapitulates matrix-vector multiplication (up to the error induced by sparsitfication). For these examples, we chose the sparsification parameter to bound the 𝓁_1_ error by 10^−8^, and so in this regime, the error we see if ≤ 10^−8^. As the population size increases, we begin combining more than two rows into any single representative row, and consequently we incur errors on the order of 10^−5^. The explanation for the increasing accuracy as *N* gets large is that our theory applies for *any* **v**, and as such is a “worst-case” bound. In contrast, our benchmarks use random **v**, and thus approximate an average case. Intuitively, our approximation results in the mean of the distribution corresponding to each row of the transition matrix being off by a little bit, and our theory bounds how off any single row can be. Yet, the mean of the distribution corresponding to some rows will be slightly too large and for others it will be slightly too small. If **v** has similar entries for a row whose mean is too large and a row whose mean is too small, some of the resulting errors will cancel. When **v** is random, as *N* gets large, more rows get grouped into a single representative row, and so there are more chances for the rows with means that are too large and too small to cancel each other out. As we will show below, in realistic scenarios (i.e., computing likelihoods) our algorithm is actually even more accurate than suggested by Figure 3. This is because likelihoods tend to be very smooth across frequencies, resulting in cancellation of errors.

Our theoretical guarantees only hold for a single matrix-vector multiplication. In theory, an approximation that is very good for a single step can become essentially arbitrarily bad over multiple rounds of matrix-vector multiplication. As such, we numerically explored the long-term accuracy of our approximation by computing transition mass functions (TMFs) — the probability of observing a given allele frequency at a particular point in the future given some current frequency. The TMF can be computed by repeatedly multiplying a vector with all zeros except for a one at the entry corresponding to the present day frequency with the single generation transition matrix. Our theory guarantees that our fast matrix-vector multiplication algorithm will result in a highly accurate approximation to the true TMF for a small number of generations, but our theory cannot determine whether the approximation gets worse over time.

To explore this, we considered a neutral model with bidirectional mutation at a rate of 1.25 ×10^−8^, and a population size of 2000, and computed transition mass functions for an initial frequency of 10%. To assess the accuracy of our approximation, we considered both the total variation distance and the symmetrized Kullback-Leibler (KL) divergence between the approximate and true TMFs. The KL divergence between two probability mass functions *p* and *q* is

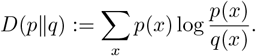

This divergence is asymmetric, so we consider the symmetrized version *D*(p‖q) + *D*(q‖p). The KL divergence is zero if and only if the two mass functions are identical, and small values indicate that the distributions are “close” in an information theoretic sense. Very roughly speaking, one over the KL divergence is approximately the number of independent observations one would need in order to distinguish the two distributions. The results are presented in Figure 4**A**, where we show that even at long time scales, our approximation remains extremely accurate. We show example transition mass functions for 10, 100, and 1000 generations in the future, where the approximation is visually indistinguishable from the true transition mass function (Figure 4**B**).

**Figure 4:**
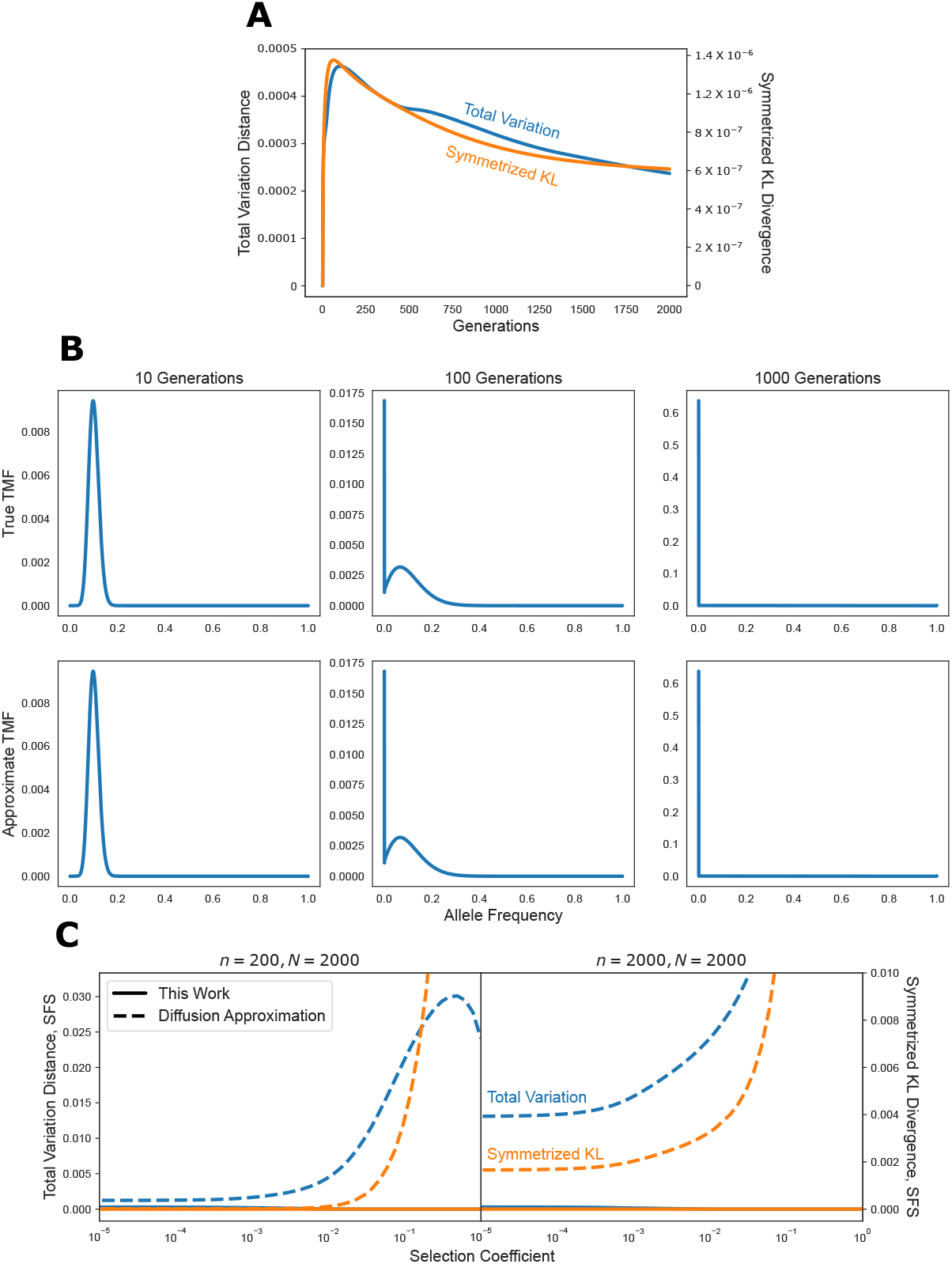
Accuracy over multiple matrix-vector multiplications. **A** Accuracy of the approximate algorithm for computing transition mass functions (TMFs) over multiple generations in terms of total variation distance (left axis label) or symmetrized KL divergence (right axis label). **B** Example TMFs at 10, 100, and 1000 generations. The first row represents the exact TMFs under the DTWF model, while the second row is the approximation derived in this work. The difference between the approximate and exact TMFs is visually indistinguishable, consistent with the low total variation distance and KL divergence. **C** Accuracy of the approximate algorithm for computing normalized site frequency spectra (SFS), as well as the commonly used diffusion approximation.

Going even further, we can consider whether we are able to recover the long-term equilibrium of the DTWF process using our approximation. To investigate this, we computed site frequency spectra (SFSs) under the DTWF model. We define the SFS more precisely in Appendix B.3, but briefly, the SFS arises in the infinite sites mutation model, which approximates the case where each position in the genome has a very low mutation rate (and hence is very unlikely to be segregating) but there are many positions across the genome, so one still expects to find some segregating sites. In this regime, mutations can only happen once per site, and so it is possible to distinguish the ancestral allele from the derived allele. The SFS is then a vector where the *i*^th^ entry is the number of positions in the genome at which there are *i* derived alleles, where *i* ranges between 1 and *n* − 1, inclusive, with *n* being the sample size. Here we consider the normalized SFS, where this vector is normalized to sum to 1, which can be interpreted as the distribution over the number of derived alleles at a randomly chosen segregating site. The normalized SFS is commonly used for demographic inference and the inference of selection coefficients [5, 6, 33].

Computing the SFS requires finding the equilibrium of a particular system, and hence can be thought of as the limit of taking infinitely many matrix-vector products. As a result, our theory on the accuracy of our approximation does not apply, and so we explored the accuracy numerically.

To compute equilibria using our approximation, we view all of the frequencies corresponding to a given representative success probability as a single “meta-state”. We then build the Markov transition matrix on these meta-states implied by our approximation to the DTWF process. Finding the equilibrium of the Markov chain on the meta-states involves solving a matrix equation with the 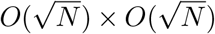 transition matrix. The resulting meta-state equilibrium is then converted to an equilibrium in the original state space by multiplying the amount of mass in each meta-state by the (truncated) Binomial PMF with that meta-state’s corresponding representative success probability. See Appendix B.2 for more details.

We compared the accuracy of our approximation to the commonly used diffusion approximation [4, 5, 18, 11], with the results presented for a range of selection coefficients and sample sizes, assuming a constant population size of 2000 in Figure 4**C**. We restricted ourselves to a population size of 2000 haploids so that we could exactly compute the ground truth by finding the equilibrium of the full DTWF transition matrix by solving a matrix equation. To compute the diffusion approximation at equilibrium we used the results presented in [34] and used as a baseline in [20].

The diffusion approximation is expected to be good when the selection coefficient is 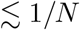 and the sample size is 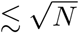 (but see [35]). In this regime, we see that both our approximation and the diffusion approximation accurately reconstruct the true normalized SFS, but with our approximation being about 4 ×more accurate in terms of total variation distance, and about 30× more accurate in terms of symmetrized KL. Yet, the diffusion approximation breaks down dramatically for large sample sizes or strong selection, while our approximation remains faithful. Indeed, for a selection coefficient of 0.01, when *n* = 200, our approximation is 250× more accurate than the diffusion approximation in terms of total variation, and 4000× more accurate in terms of symmetrized KL. Similarly, when *n* = *N* = 2000, even when the selection coefficient is zero, our approximation is about 45× more accurate in terms of total variation, and 3800× more accurate in terms of symmetrized KL. Taken together, we see that our approximation is highly accurate across the full spectrum of sample sizes and selection coefficients.

### 3.2 Impact of mutation, selection, and demography on the DTWF model

Having established the accuracy of our model, we turned to using it to better understand the use of large sequencing cohorts for estimating selection coefficients. There has recently been growing interest in using the frequency of loss-of-function variants (LoFs) in large-scale exome sequencing projects to estimate measures of gene constraint [23, 24, 25, 26, 27, 28]. LoFs are variants, such as early stop codons, splice-disrupting variants, or frameshifts, that result in the gene failing to make a viable protein. To a first approximation, all of the LoFs within a gene have roughly the same strength of selection acting against them, as they all have similar effects on the protein. As such, LoFs are attractive for studying selection as we can pool information across all of the LoF variants within a gene to estimate a single LoF selection coefficient for that gene.

Previous approaches have relied on deterministic approximations [24], simulations [23, 28], or *ad hoc* methods and models [25, 26, 27] to infer selection coefficients from LoF data. These approaches have yielded widely-used measures of gene constraint and important insights into the landscape of constraint on human genes. Yet, without more principled computational machinery for computing likelihoods, it can be difficult to estimate the gains in power we might expect to see in different datasets. For example, how does increasing the sample size affect our power to estimate selection coefficients? How does demography affect power? Does sampling from a population that has experienced recent growth affect power? What about a recent bottleneck? How does recurrent mutation affect power? Are some types of variants more informative than others? In this section we use our machinery to answer these questions.

To understand how sample size, mutation rate, and demography interact to affect power for estimating selection coefficients, we considered a variety of each of these parameters. In particular, we considered sample sizes ranging from *n* = 10 diploids to *n* = 300,000 diploids, encompassing the range from small pilot studies in non-model organisms to biobank-scale datasets. To understand the impact of mutation rates and recurrent mutations, we considered a low mutation rate typical of transversions in humans (2.44× 10^−9^ per generation) as well as a high mutation rate typical of methylated CpG sites in humans (1.25×10^−7^ per generation) [25]. For demographies, we considered slight modifications of two demographies estimated using MSMC [7] applied to the 1000 Genomes Project [36] — one demography estimated from individuals labeled by the 1000 Genomes Project as “Utah residents with Northern and Western European ancestry” (CEU) and one estimated from individuals labeled as “Yoruba in Ibadan, Nigera” (YRI). The CEU demography consists of a strong bottleneck corresponding to the out-of-Africa event, and recent explosive growth, whereas the YRI demography lacks a bottleneck and has remained roughly constant in size over time (Appendix Figure A1). See Appendix E for more details.

We used these different sets of mutation rates, sample sizes, and demographies in a DTWF model. Specifically, following previous work we focused on a diploid model of additive selection on LoFs [23, 24, 28], where having one copy of the LoF variant results in a fitness reduction of *s*_*het*_, while having two copies results in a fitness reduction of *s*_*hom*_ := 2*s*_*het*_ (but with fitness lower bounded by 0 in the event that *s*_*het*_ > 0.5). Our computational machinery was developed for haploid populations and only tracks allele frequencies and not genotype frequencies. To approximate the diploid model of selection we set the expected frequency in the next generation, *p*(*f*), as

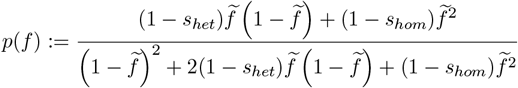

with

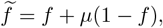

where *f* is the frequency of the LoF in the current generation, so that 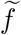 is the frequency following mutation at rate *μ*. Under strong selection frequencies will generally be low, so we ignore back-mutation. This model matches the expected frequency change under the diploid selection model assuming Hardy-Weinberg equilibrium [1].

Without back-mutation, the equilibrium of this process is the degenerate state where the population is fixed for the LoF mutation, which is obviously not biologically realistic. One could instead turn to the commonly-used infinite sites model, but this comes with two issues. First, any particular site must be non-segregating with probability one as the infinite sites model assumes an infinitesimally small per-site mutation rate balanced by an infinitely large mutational target size. This assumption may be realistic when considering mutations genome-wide, but certainly breaks down when looking at single LoFs, or even across LoFs within a single gene.

Second, since the mutation rate per site is infinitesimally small under the infinite sites approximation, the probability of recurrent mutation is also 0. Recurrent mutation in this context refers to the same allele being generated at the same site via independent mutation events. For small sample sizes and small mutation rates, the probability of independent mutations happening at the same site is extremely small, explaining the popularity of the infinite sites model. Yet, for the CpG mutation rate we know that recurrent mutations are common and play an important role in shaping diversity [37].

Instead of relying on the infinite sites approximation, our computational machinery allows us to easily condition the evolutionary dynamics of an allele at a single site on non-fixation. Essentially, when a new LoF allele enters the population, we ignore any scenarios where it drifts to fixation in the population. Looking backward in time, in a finite population alleles must at some point in the past either have been fixed or totally absent from the population. Since we are explicitly not allowing the LoF allele to have been fixed at any point in the past, there must have been some point in the past at which the LoF arose as a new mutation in a population monomorphic for the non-LoF allele. In this way, there is a well-defined notion of an ancestral allele and a derived allele. The DTWF model conditioned on non-fixation is well-behaved and has a non-trivial equilibrium. Additionally, it allows us to easily model recurrent mutations and obtain a non-zero probability of an individual site being segregating. See Appendix B.3 and B.4 for more details surrounding this subtlety.

Before investigating the impact of selection, we wanted to see if modeling recurrent mutations was necessary for biobank-scale datasets under our models. Recently, analytical results for recurrent mutations in the coalescent (i.e., in the diffusion limit, and assuming neutrality) have been developed [38]. Here we also focus on the neutral case for simplicity, and our results our qualitatively similar to those in [38]. The approaches are complimentary: the results in [38] are analytic, while ours must be obtained numerically. For our machinery it is no more difficult to consider cases with various types of selection, whereas obtaining coalescent-based results in the presence of selection would be difficult.

We considered something analogous to the SFS under our model — the probability of observing a given frequency of a derived allele conditioned on the site being segregating. We show the results for the CEU demography in Figure 5, where we see that for large sample sizes recurrent mutations have a large effect on the frequency spectrum, with singletons being almost half as likely under the CpG mutation rate compared to the transversion mutation rate. This is somewhat counterintuitive — one might expect that under a high mutation rate there would be more rare variation, and that *is* true in absolute terms as there are more segregating sites, but given that a site is segregating, rare variants actually become less likely under higher mutation rates. Indeed, at a sample size of *n* = 300,000 diploids, the probability that a CpG is segregating is 0.678, while for transversions it is only 0.022. At smaller sample sizes the impact of recurrent mutation is negligible for realistic mutation rates, with the probability that a site is segregating being 0.033 for CpGs and 0.0007 for transversions for a sample size of *n* = 100 diploids. The results for the YRI demography are qualitatively consistent (Figure A2), although some of our modeling choices result in an unusual and interesting non-convex frequency spectrum, which we discuss in detail in Appendix E.

**Figure 5:**
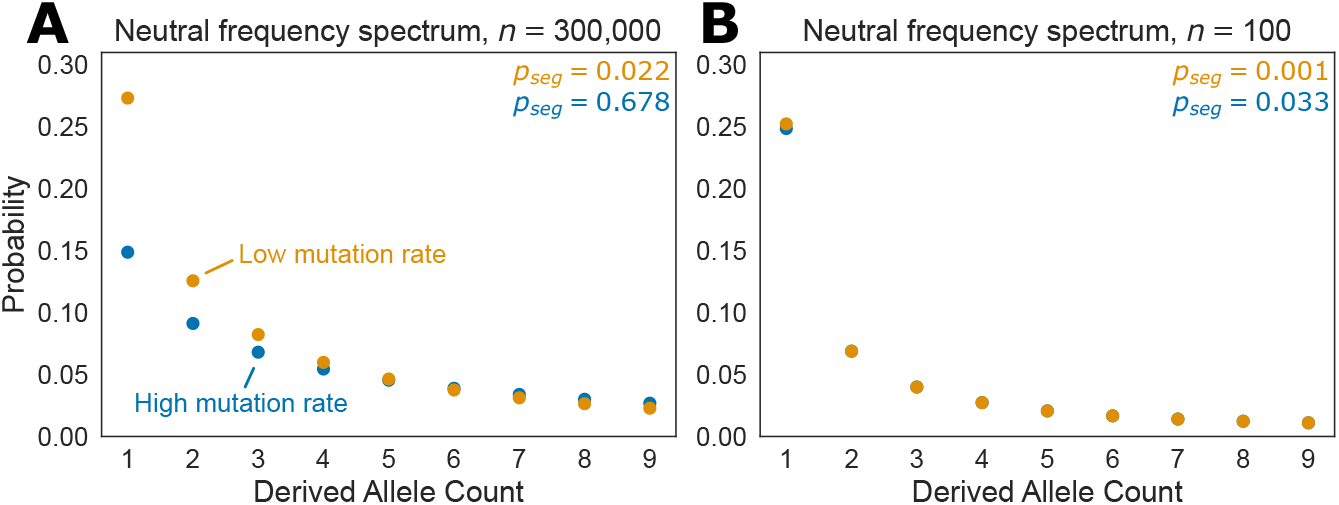
Frequency spectra under the CEU demography for sample sizes *n* = 300,000 diploids (**A**) and *n* = 100 diploids (**B**) for a low mutation rate (2.44×10^−9^ per generation) and a high mutation rate (1.25×10^−7^ per generation). The low mutation roughly corresponds to the rate of a transversion in humans, while the high mutation rate rough corresponds to the rate of mutation at a methylated CpG. The probability of a site being segregating is reported as *p*_*seg*_.

We next turned to understanding the impact of mutation rates on using LoF frequencies to estimate *s*_*het*_.

To this end, we computed the likelihood of observing each possible frequency (including 0) in a given sample for a range of values of *s*_*het*_ ranging from well below the nearly neutral limit (*s*_*het*_ = 10^−6^) all the way up to nearly lethal (*s*_*het*_≈ 1). We show the results for the two mutation rates we considered for a sample of size 300,000 diploids from the CEU demography in Figure 6. The results show that rare variants are weakly indicative of strong selection, but otherwise observing an LoF of a given frequency acts as a soft threshold on *s*_*het*_. For example, a doubleton confidently rules out *s*_*het*_ > 0.1, but is otherwise essentially equally consistent with any value of *s*_*het*_. Similarly, an LoF at 1% frequency rules out any *s*_*het*_ > 0.002, but is otherwise relatively uninformative. The results are qualitatively similar across the two mutation rates, but very low frequency or non-segregating CpGs are much more evidence in favor of strong selection than non-segregating transversions, consistent with recent work by Agarwal and Przeeworski showing empirically and via simulation that a non-segregating CpG at similar sample sizes is enough to confidently reject neutrality [39].

**Figure 6:**
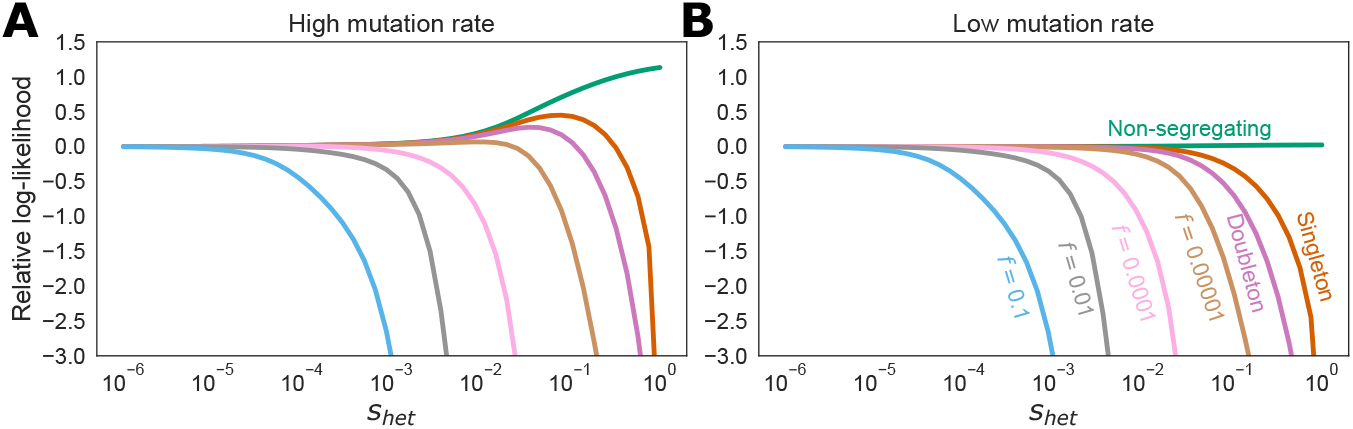
Likelihoods for a sample of size *n* = 300,000 diploids from the CEU demography assuming either a high mutation rate of 1.25 *×* 10^−7^ per generation (**A**) or a low mutation rate of 2.44 × 10^−9^ per generation (**B**).

To more precisely quantify how informative different sample sizes, datasets, or mutation rates are for estimating selection we used the Fisher Information, ℐ. Fisher Information quantifies the expected curvature of the likelihood function at a given value of *s*_*het*_ and can be thought of as an effective sample size multiplier in terms of number of variants. In the DTWF model, information is additive across independent sites, so a setting with twice the Fisher Information would require half as many independent variants to achieve the same level of accuracy roughly speaking. More formally, the Cramer-Rao lower bound from statistics shows that any unbiased estimator of *s*_*het*_ must have variance greater than 1*/ℐ*. As such, the Fisher Information can be thought of as being inversely related to the variance of the best unbiased estimator of *s*_*het*_. In our setting, the Fisher Information is defined as

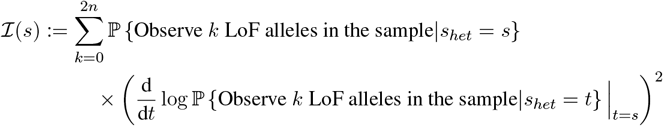

which we can compute using the likelihood curves shown in Figure 6 via numerical differentiation. Note that the Fisher Information depends on the parameterization of *s*_*het*_, and here we compute the Fisher Information for log10 *s*_*het*_ to match the parameterization shown in Figure 6. As a result, the Fisher Information can be thought of as being related to how many orders of magnitude the uncertainty in *s*_*het*_ should span.

We began by investigating the information content of CpGs and transversions for estimating *s*_*het*_. For both the CEU and YRI demographies, and across all sample sizes we find that the information content of a CpG is always between 49× and 51.5× greater than that of a transversion. This makes intuitive sense as the mutation rate is about 51.2× higher for CpGs, indicating that we would expect CpGs to be segregating roughly 50× as often as transversions. CpGs are usually slightly less than 51.2× as informative as transversions across different values of *s*_*het*_, indicating that there are some diminishing returns.

Next we turned to how information grows with sample size. Unlike standard statistical settings, sampling additional individuals does not provide completely independent information, and one might expect information to plateau as the sample size grows. Indeed, since individuals share a common genealogy, as additional individuals are added to a sample they are increasingly likely to be closely related to someone already in the sample, and hence provide little additional information. This is indeed borne out in our results (Figure 7**A**), where we see that increasing sample sizes provide diminishing returns in terms of information. Yet, this effect is not uniform across the space of *s*_*het*_ values. Our results suggest that increasing sample sizes beyond the approximately 140,000 individuals in gnomAD [25] only provides additional information for the most extremely selected variants (*s*_*het*_ > 0.02). This highlights that there is a fundamental limit on how much we can hope to learn about the selective pressure on genes from LoF data alone — at current sample sizes we have already saturated the amount of information we might hope to obtain for a wide range of selection coefficients. Further increases in sample sizes will only help resolve the selection coefficients for the genes with the most extreme effects on fitness.

**Figure 7:**
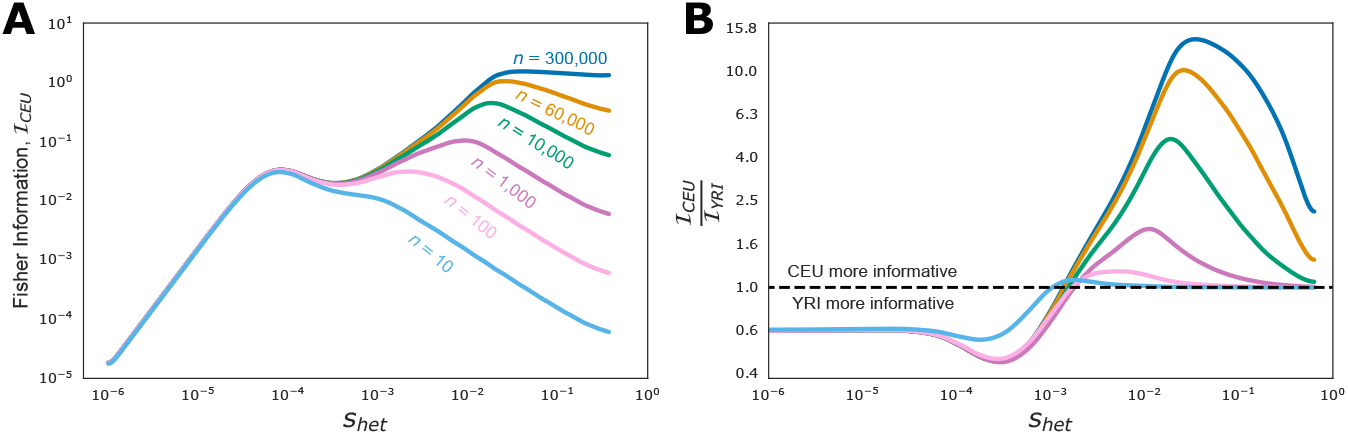
**A** Fisher Information as a function of *s*_*het*_ in samples from the CEU demography of different sample sizes. **B** Fisher Information for a sample from the CEU demography relative to the Fisher Information for a sample of the same size from the YRI demography. Points above the dashed line indicate settings where the CEU sample provides more information, points from below the dashed line indicate settings where the YRI sample provides more information.

Finally, existing exome sequencing cohorts consist primarily of individuals that are genetically similar to the CEU individuals in the 1000 Genomes Project [25, 40], raising questions of whether we might be able to better estimate selection by looking at samples of individuals with other genetic ancestries. For example, is it more informative to have a sample of a given size from a population that underwent the CEU demography or a population that underwent the YRI demography? Interestingly, we find that the answer depends strongly on *s*_*het*_ and the sample size. For the smallest sample sizes (e.g., *n* < 100 diploids) samples from the YRI demography or the CEU demography are comparably informative for selection coefficients above 0.001, but samples from the YRI demography are almost twice as informative for selection coefficients below 0.001 (Figure 7**B**). The dominance of a sample from the YRI demography for low selection coefficients remains across sample sizes, but as sample sizes increase, samples from the CEU demography become increasingly more informative for large selection coefficients. For example, at a sample size of *n* = 1,000 diploids, a sample from the CEU demography is nearly twice as informative for an *s*_*het*_ of 0.01, and for a sample of size *n* = 300,000 a sample from the CEU demography is about 15 times as informative for an *s*_*het*_ of 0.1.

The relative Fisher Informations for samples from the CEU and YRI demographies can be understood in terms of those demographies. Variants under weak selection are older, and due to the out-of-Africa bottleneck, those variants will have experienced stronger drift under the CEU demography than the YRI demography. Hence, samples from the CEU demography contain more “demographic noise” due to drift for such variants. Conversely, the recent explosion in population size in the CEU results in the opposite phenomenon for strongly selected variants which likely arose more recently, resulting in more information for large values of *s*_*het*_. Finally, for small sample sizes there is little power to get at the rare, recent variants indicative of strong selection, regardless of demography, explaining why samples from the CEU demography only become more powerful than samples from the YRI demography for strong selection at large sizes.

## 4 Discussion

Here we presented an approach to approximate the transition matrix of the DTWF process that is provably accurate and allows us to compute likelihoods in *O*(*N*) time. We showed that our approach can scale to population sizes in the billions, and is highly accurate. Our approach relied on two key observations: the transition matrix of the DTWF process is approximately sparse, with only 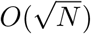 entries contributing appreciably to the mass in each row, and the matrix is approximately low rank, where the matrix can be replaced by one with only 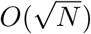 unique rows while incurring a small error.

We used our approach to understand how increasing sample sizes will help estimate the strength of selection acting against gene loss-of-function. We found that increasing sample sizes beyond those currently available will only provide additional information for the most strongly selected genes. For genes with anything weaker than the most extreme selection, current samples provide essentially as much information as can ever be obtained from LoF data from individuals closely related to the 1000 Genomes CEU sample.

Our approach may seem similar to choosing a discretization scheme for the PDE that describes the WF diffusion, but the approaches are distinct. The PDE discretization approach starts with the DTWF model, passes to a continuum limit to obtain a PDE, and then discretizes *that* continuous process. Yet, the WF diffusion is only valid for fixed frequencies as *N*→ ∞, indicating that in practice the continuous process is not a good approximation for frequencies close to 0 or 1. As such, any discretization of the WF diffusion must also be inaccurate near the boundaries. Instead, here we propose directly coarse graining the underlying discrete process without passing to a continuum limit.

In some ways, our approach is reminiscent of the scaling approach used in simulations [41]. In forward-intime simulations, it can be computationally onerous to simulate a large population. To avoid this, one chooses a scaling factor, such as 10, and simulates a population 10× smaller, but increases the mutation rates, recombination rates, and strengths of selection by a factor of 10. Additionally, each generation in this scaled model counts for 10 generations in the unscaled model. This scaling is chosen so that the rescaled population converges to the same WF diffusion as the original process. Yet, this rescaling is only trustworthy for frequencies ≫1*/N*. Here, we do not rescale parameters, but we do group states into “meta-states”, and we group states more aggressively when the frequency is close to 0.5, and less aggressively for frequencies near 0 or 1. Whether a similar idea of frequency-adaptive rescaling could be incorporated into simulation to improve speed while remaining accurate is an interesting area for future research.

Another view of our method is that we are replacing a difficult set of transition distributions with a simpler set. This idea is very general and different approaches could be taken. For example, it may be possible to match the first several moments using only a very small number of non-zero entries. Such extremely sparse transition matrices could result in highly accurate and very computationally efficient approximations. The approach presented here is just one possibility in this vein, and exploring alternatives could be a fruitful direction for future research.

Our results are quite general, and can be readily extended to multiple alleles, multiple populations, or multiple loci. All of these can be treated as processes defined by a transition matrix of sub-Gaussian probability mass functions, and similar arguments to those used here can be applied to show that such transition matrices have approximately sparse rows, and are approximately low rank. These arguments should result in comparable speedups, but unfortunately, this direct approach of computing likelihoods using the forward transition matrix necessarily comes with a steep computational cost in these settings. For example, simply to list all of the possible configurations of a population of *N* individuals at two biallelic loci requires *O*(*N* ^3^) time [42]. To list all of the possible configurations for 3 loci requires *O*(*N* ^7^) time, and in general *k* loci requires 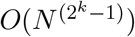 time. There may be additional approximations that can be made in these cases, but simply approximating the transition matrix as we do here will not be enough to handle these more combinatorially difficult cases.

Throughout, we have assumed that the goal is to approximate the underlying DTWF model while maintaining computational efficiency. In general, however, no population will exactly follow any simple DTWF model — in many populations there will be fine-scale geographic population structure [43], assortative mating [44], overlapping generations, and so on. While these complications may make the DTWF model seem overly simplistic, the WF diffusion must be an even worse approximation as it also implies unrealistic family size distributions for large sample sizes [35]. In any case, our approach may also be useful for more complex models (e.g., general Cannings’ exchangeable models [45, 2]), as long as transitions have the two properties of being restricted (with high probability) to a small subset of the state space, and transition probability mass functions for nearby states being similar enough to be nearly indistinguishable. The extent to which these two properties are true will determine the extent of the speedup offered by our approach, and will depend on details of the underlying model. For example, the forward-in-time models that result in coalescents with multiple mergers [46, 47] or simultaneous multiple mergers [48, 49] often correspond to “sweepstakes reproduction” where a single individual may spawn a sizable fraction of the next generation. Under these models, a large sweepstakes reproduction event could cause an allele to dramatically change frequency in a single generation indicating the the transition density for any state is *not* approximately sparse, and the approach used in this paper would not result in a large speedup.

Here we focused on the problem of computing the likelihood of observing a given number of derived alleles at present, but our speedups apply to time series data as well, which is frequently encountered in ancient DNA. Several methods have been developed that treat the true allele frequency at a given time as a hidden state in a hidden Markov model (HMM). This frequency then evolves through time according to the transition matrix of either the DTWF [50], WF diffusion [51], or some other approximation [52], with sampled genotypes as the observations in the HMM. These HMMs have been particularly useful in estimating the strength of natural selection acting on individual loci, and our results can be used in these methods to speed up computations while directly approximating the DTWF model.

Our implementation is in `pytorch` [53], which allows for backward mode automatic differentiation, enabling the computation of gradients of functions of the likelihood with respect to selection coefficients or mutation rates. Unfortunately, backward mode automatic differentiation requires storing the entire “computation graph” in memory. In our setting, this corresponds to storing representations of the approximate transition matrices at each generation, which may become memory intensive in models where the non-equilibrium portion spans many generations. Indeed, throughout this paper we resorted to using numerical approximations to the gradient to avoid these issues. Since our likelihood computation essentially just involves repeated matrix-vector multiplication, one may view it as a very deep neural network with linear activations, and backward mode automatic differentiation proves to be memory intensive in those applications as well [54]. Our approach is also mathematically similar to using discretization to integrate a linear ODE forward in time, another application which essentially boils down to repeated matrix-vector multiplication. In that setting powerful methods have been developed which essentially solve the ODE forward to calculate the likelihood and then backward to obtain gradients, which avoids the need to store the computation graph in memory [55]. Extending this approach to our setting is a promising approach to obtain gradients without resorting to numerical approximation.

As modern datasets approach sample sizes of hundreds of thousands to millions, new scalable approaches are needed in population genetics. This onslaught of data is a blessing, but more work like this — developing provably accurate, scalable approaches — is needed to keep up and allow us to extract useful insights from these ever growing sample sizes. Yet, care should be taken as our results show that larger sample sizes are not always helpful. For the problem of estimating selection coefficients, larger sample sizes will never provide less information, but for many genes they will not provide more information.

## Acknowledgements

We would like to thank Ian Holmes for discussions about coarse graining Markov chains and Paul Jenkins for discussions about conditioning on non-fixation. We would like to thank Clemens Weiß and Amy Ko for helpful discussions and valuable feedback on the manuscript. This work was supported by NIH grant R01HG011432.

## A Formal Theoretical Results and Proofs

### A.1 Notation

Throughout we will use the notation *B*_*N,p*_ for a Binomial distribution with sample size *N* and success probability *p*. We will also restrict our attention to what we define as *ordered Binomial transition matrices*, of which many DTWF models are special cases. We say that a matrix, 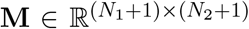 is a *Binomial transition matrix* if each row of **M** is the probability mass function of a Binomial distribution. That is, for each *i* the *i*^th^ row of **M** is the probability mass function corresponding to a 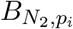 distribution. Furthermore, we say that **M** is an *ordered* Binomial transition matrix if it is a Binomial transition matrix with the further property that the rows are ordered by success probability; that is, *p*_0_ ≤ *p*_1_ ≤ … ≤ *p*_*N*_1.

### A.2 Construction of approximate transition matrix

The main goal of this section is to prove that we can accurately approximate any Binomial transition matrix with a highly structured matrix for which matrix-vector multiplication is much faster than standard. The overall crux of this proof is twofold.

First, nearby rows in a Binomial transition matrix are close in total variation distance, indicating that we can replace one row with a copy of a nearby row while only incurring a small error. To prove this, we will show that the total variation distance between *B*_*N,p*_ and *B*_*N,p*_ *′* is small <u>so</u> long as *p* and *p*′ are close in a particular sense. This will allows us to partition the interval [0, 1] into 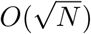 blocks such that for any *p* and *p*′ in the same block the corresponding Binomial distributions are guaranteed to be close in total variation distance. This in turn shows that *any* Binomial transition matrix for a population of size *N* can be replaced by a matrix with only 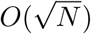 unique rows while controlling the row-wise error.

Second, we will use a classical result to show that the tails of the Binomial distribution are incredibly light — it is very unlikely to sample a value far away from the mean of a Binomial distribution, and in particular, one incurs only a small approximation error by only considering the possibility of sampling something within 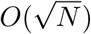 of the mean. This will allow us to replace each row of the Binomial transition matrix by a sparse vector with only 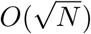 nonzero entries, while only incurring a small approximation error.

To start showing that we can partition [0, 1] into 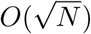 blocks where Binomial distributions with success probabilities in the same block have bounded total variation distance, we begin with the block that includes 0.

#### Lemma 1.

*For any p* ≤ 1 − (1 − *ε*)^1*/N*^,

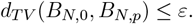

*In particular, for fixed ε, p can be as large as* 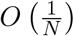 *while maintaining a total variation distance to a Binomial distribution with success probability* 0 *at most ε*.

*Proof*. Let *X* ∼ *B*_*N*,0_ and *Y* ∼ *B*_*N,p*_. Note that *X* must be 0 with probability 1, and cannot take any other value. The total variation is then, by definition,

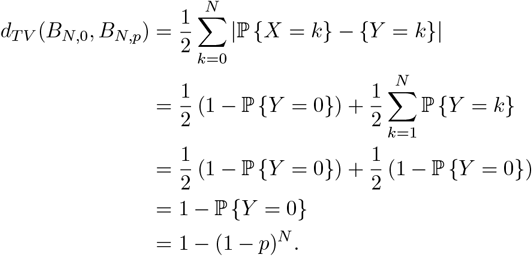

This is obviously monotonically increasing in *p*, and solving for *p* we obtain that

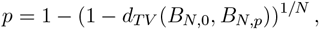

so to obtain a total variation distance at most *ε* we need

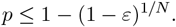

Finally, rewriting (1 − *ε*)^1*/N*^ as 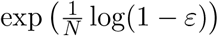 and using the convergent series expansion, we see

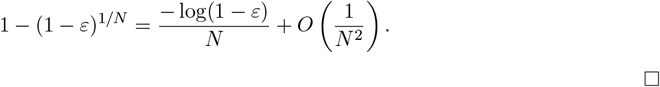

The following lemma, stated in [56] and proved in [57] bounds the total variation distance between Binomial distributions with success probabilities away from 0 and 1.

#### Lemma 2.

*([56, Equations (2*.*15) and (2*.*16)]) For any p* ∈ (0, 1) *and any* δ ∈ (0, 1 − *p*),

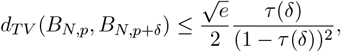

*where*

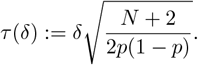

For our purposes, we just need to know how far away *p*′ can be from *p* before incurring an unacceptable total variation distance, which we can obtain by loosening the above bound and rearranging.

#### Lemma 3.

*Fo*l*r any p* ∈ (0, 1*/*2), *there exists a constant c*_*ε*_, *that depends on ε but not on p or N such that for any* 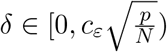 *we have*

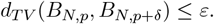

*Proof*. First, note that for any *p* ∈ (0, 1*/*2) and *N* ≥ 1 we have

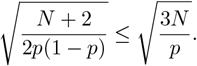

Letting 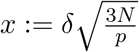, we have by Lemma 2 that

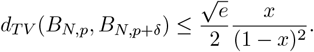

The right hand side is obviously monotonically increasing in *x* on [0, 1) from a value of 0 at *x* = 0 to infinity as *x* approaches 1. Furthermore, the equation does not contain *p* or *N* (except in the definition of *x*). Therefore, there exists a 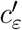 independent of *p* and *N* such that when 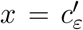 the right hand side is *ε*, and hence for any 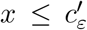 we have that *d*_*T V*_ (*B*_*N,p*_, *B*_*N,p*+δ_) ≤ *ε*. Using our definition of *x* and solving for δ completes the proof.

With these Lemmas in place, we can now prove our result on partitioning [0, 1] such that Binomial distributions with success probabilities in the same block have bounded total variation distance.

#### Lemma 4.

*For fixed ε, there exist* 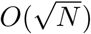 *breakpoints* 0 = *p*_0_ *< p*_1_ *< p*_2_ *< … < p*_*K*_ = 1 *such that for any*

*p and p*′ *within in the same interval (i*.*e*., *there exists an i such that p, p*′ ∈ [*p*_*i*_, *p*_*i*+1_]*) we have*

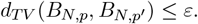

*Proof*. Note that by symmetry, we only need to consider partitioning the space [0, 1*/*2] using 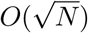 break-points. By Lemma 3 we can control the total variation distance of all distributions between a breakpoint *p*_*k*_ and *p*_*k*+1_ by taking

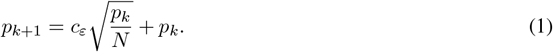

Therefore we have that the total variation distance is less than *ε* between any Binomials of size *N* with success probabilities between *p*_*k*_ and *p*_*k*+1_.

Now, we will prove by induction that there exists a constant, *α*_*ε*_, that depends on *ε* but not on *N*, such that

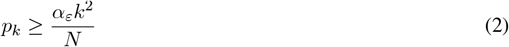

The base case of *p*_1_ is handled by Lemma 1. Now, suppose that Equation 2 holds for *p*_*k*_, then, by Equation 1 and the inductive hypothesis,

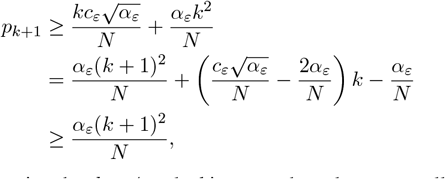

where the last line follows by noting that *k* ≧ 1 and taking *α*_*ε*_ to be at least as small as 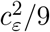.

This proves Equation 2. Then, to partition [0, 1*/*2] we can compute breakpoints using the recursion in Equation 1 until we reach the first breakpoint larger than 1*/*2. By Equation 2 we need at most

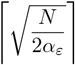

breakpoints, which is 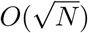, to partition the space, completing the proof.

We now turn to the task of showing that for a given Binomial distribution almost all of the mass is on outcomes within 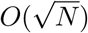 of the mean. This result follows straightforwardly from Hoeffding’s celebrated inequality [30], which we include for completeness

#### Lemma 5.

*(Hoeffding’s Inequality [30, Theorem 1]) Let X* ∼ *B*_*N,p*_, *then for k* ≤ *Np*,

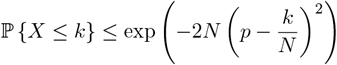

*and for k* ≥ *Np*,

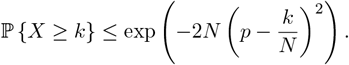

#### Lemma 6.

*Let*

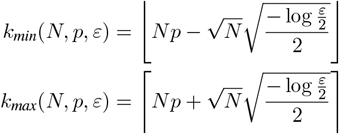

*and define* 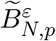 *as the distribution obtained by conditioning B*_*N,p*_ *to take values in*

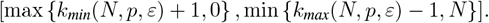

*Then*,

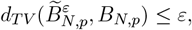

*and the mass function for* 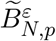 *contains only* 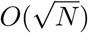 *non-zero entries*.

*Proof*. That the mass function of 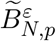 contains only 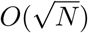 non-zero entries is obvious from its construction. To bound the total variation distance, let *X* ∼ *B*_*N,p*_, and let 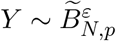 By construction,

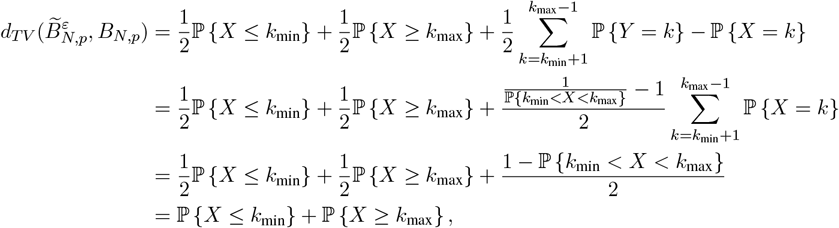

where we dropped the dependence of *k*_min_ and *k*_max_ on *N, p*, and *ε* for notational convenience. Then, by Lemma 5,

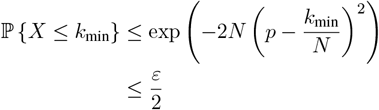

where the second line follows from our choice of *k*_min_. An analogous computation shows that

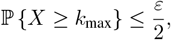

completing the proof.

The following lemma will be used to show that as long as a Binomial transition matrix is ordered then we can assign the success probabilities for each of its rows to a set of blocks in linear time.

#### Lemma 7.

*Consider* 0 ≤ *v*_1_ ≤ *v*_2_ ≤ … ≤ *v*_*N*_ ≤ 1, *and a partition of the space* [0, 1] *defined by breakpoints* 0 = *p*_0_ *< p*_1_ *<* … *< p*_*K*−1_ *< p*_*K*_ = 1. We may compute index sets 𝒮_1_, …, 𝒮_*K*_ *in O*(*N* + *K*) *time such that for all k for each i* ∈ 𝒮_*k*_ *we have that p*_*k*−1_ ≤ *v*_*i*_ ≤ *p*_*k*_, with 𝒮_1_ ∪ *…* ∪ 𝒮_*N*_ = *{*1, …, *N*} *and* 𝒮_*i*_ ∩ 𝒮_*j*_ = ∅ *for all i ≠j*.

*Proof*. We begin with *v*_1_ and we find the first *k* such that *p*_*k*_ that is at least as large as *v*_1_. We may then search from *v*_2_ onward until we find the first *j* such that *v*_*j*_ is larger than *p*_*k*_. If no such *j* exists, then set *j* to be *N* + 1. We assign 1, …, *j* − 1 to index set 𝒮_*k*_. This is a valid assignment as we have *p*_*k*−1_ ≤ *v*_1_ ≤ … ≤ *v*_*j*−1_ ≤ *p*_*k*_. We may then repeat this process —we next find the first *k*_2_ such that 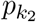 is at least as large as *v*_*j*_, and we find the first *j*_2_ such that *v*_*j*_2 is larger than 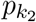 (or if such a *j*_2_ does not exist, set *j*_2_ to *N* + 1), assigning *j*, …, *j*_2_ − 1 to 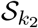. We can repeat this process until all of the *v*’s have been assigned to an index set. Since both the *v*’s and *p*’s are sorted we can do these searches starting where the previous search left off, and once we reach the end of both the *v*’s and the *p*’s we have assigned all of the *v*’s to an index set. Thus, we only have to consider each *v* and *p O*(1) times, resulting in a total runtime of *O*(*N* + *K*).

Combining the previous lemmas, we can construct a highly structured matrix that accurately approximates an ordered Binomial transition matrix in *O*(*N*) time.

#### Proposition 1.

*Let* 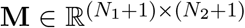 *be an ordered Binomial transition matrix. We can build a representation of a matrix* 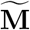 *with the following properties:*

- 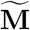 *has* 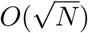 *unique rows*,
- *Each row of* 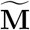 *has at most* 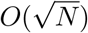 *non-zero elements*,
- 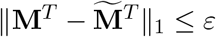

*Furthermore, we can construct this representation in O*(*N*_1_ + *N*_2_) *time and store it in O*(*N*_1_ + *N*_2_) *space*.

*Proof*. Since **M** is an ordered Binomial transition matrix, each row is a probability mass function of the form 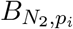 with nondecreasing *p*_*i*_. By Lemma 4 we can partition the space [0, 1] into 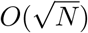 blocks such that for any *p*_*i*_, *p*_*j*_ in the same bucket the total variation distance between the rows is *ε/*4. We can assign each of the Binomial probability mass functions to these blocks in 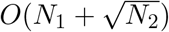 time by Lemma 7. Each row of the matrix with an index in the same index set can then be replaced by a Binomial probability mass function with an arbitrary “representative” success probability contained in that block. Since each row is getting replaced by a row corresponding to a distribution with total variation distance less than *ε/*4, we have that the 𝓁_1_ distance between each row is less than *ε/*2. Finally, by Lemma 6 we can replace the distribution for each representative row, by an 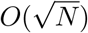 sparse version while only incurring a further total variation distance of *ε/*4, resulting in a further 𝓁_1_ distance of *ε/*2. By the triangle inequality, each final row is then at most *ε* away from the original row in 𝓁_1_ distance.

Since there are only 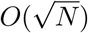 unique rows and each each of these have only 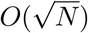 elements, once we have assigned rows to their representative rowss constructing this sparse representation only requires *O*(*N*_2_) time, resulting in an overall runtime of *O*(*N*_1_ + *N*_2_). Representing 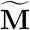 requires space, as storing the 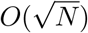 unique rows, each with 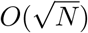 non-zero entries requires *O*(*N*_2_) space, and then we must also store an index for each row in 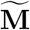 indicating which representative row to use, requiring *O*(*N*_1_) space.

The requirement in Proposition 1 that the Binomial transition matrix be ordered is so that we can determine which rows can be replaced with which representative rows in linear time. In the absence of such information (i.e., if the Binomial success probabilities for each row of the matrix are arbitrary) then we need *O*(*N* log *N*) time to determine the representative rows to use for each row.

The following lemma shows that matrix-vector products can be made substantially faster for matrices with a limited number of unique rows where each of those rows are sparse.

#### Lemma 8.

*Let* **M** ∈ ℝ^*N ×P*^ *be a matrix with n unique rows, where each row has at most s non-zero elements. Furthermore, suppose we have index sets 𝒮*_1_, …, 𝒮_*n*_, *with 𝒮*_1_ ∪ *…* ∪ 𝒮_*n*_ = *{*1, …, *N} and* 𝒮_*i*_ ∩ 𝒮_*j*_ = ∅ *for all i* ≠ *j with the property that if rows k and* 𝓁 *are in the same index set than those rows of* **M** *are identical. Then, for any vector* **v** ∈ **R**^*N*^ *the matrix-vector product* **M**^*T*^ **v** *can be computed in O*(*N* + *P* + *ns*) *time*.

*Proof*. Let **M**_*i*_ be the *i*^th^ row of **M**. For the desired matrix product we need to compute

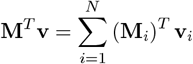

Since the index sets cover all of the indices with no overlaps, we can rewrite the above sum as:

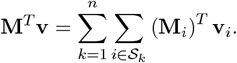

Let *i*_1_, …, *i*_*n*_ be arbitrarily chosen elements from each of the *n* index sets. Then, by noting that if *i* is in **S**_*k*_, we have by assumption 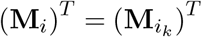, which allows us to pull the inner summation inward:

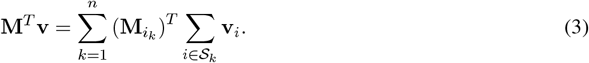

The inner sum is now a sum of scalars, so for a particular *k* it costs *O*(|*S*_*k*_|) to compute. To compute it across all *k* thus costs *O*(*N*) time. Then, since each 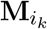 only contains *s* non-zeros, we can multiply each by a scalar, and then sum up all *n* of them in *O*(*ns* + *P*) time. To see this, note that we can initialize a “running total” vector of *P* zeros, and then for each 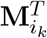 we only need to update the running total in the *s* entries at which 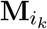 is non-zero.

Combining Proposition 1 and Lemma 8, we immediately arrive at our main result.

#### Theorem 1.

*Let* 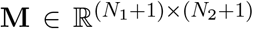 *be an ordered Binomial transition matrix. We can replace* **M** *by a matrix* 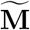 *such that* 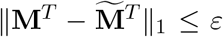, *and such that computing matrix vector products of the form* 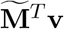 *requires O*(*N*_1_ + *N*_2_) *time*.

*Proof*. Each row of **M** is a probability mass function of the form 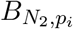 with nondecreasing *p*_*i*_. By Proposition 1 we can approximate this matrix by a matrix with at most 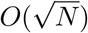 unique rows, each with at most 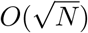 non-zero entries, while maintaining 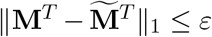, and we can construct a representation of this matrix in *O*(*N*_1_ + *N*_2_) time. By Lemma 8 we can perform matrix-vector multiplication with such a matrix in *O*(*N*_1_ + *N*_2_) time.

### A.3 Sample likelihoods

In this section, we discuss how to efficiently obtain the likelihood of observing *k A* alleles in a sample of size *n* from the vector of probabilities of observing different numbers of *A* alleles in a population of size *N*. While this may seem like a substantially different problem, we show that similar considerations to speeding up matrix-vector multiplication for Binomial transition matrices can be applied to this subsampling problem as well. In particular, it is generally assumed that the sample of size *n* is drawn without replacement from the population of size *N*. Suppose that there are *K A* alleles in the population, then the probability of drawing some number of *A* alleles in a sample of size *n* is determined by the Hypergeometric distribution with parameters *N, n*, and *K*, which we will denote in this section by *H*_*N,K,n*_. Since we do not observe the frequency of the *A* allele in the population, we should integrate over this latent variable. Thus, if **v**_sample_ ∈ ℝ^*n*+1^ is the vector of sample probabilities, and **v** ∈ ℝ^N+1^ is the vector of population probabilities, then,

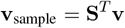

where **S** is a “sampling matrix” where the *k*^th^ row is the probability mass function corresponding to the distribution *H*_*N,k,n*_.

Surprisingly, our results about Binomial transition matrices also apply in modified forms to Hypergeometric sampling matrices. In particular, we show below that such matrices have rows that are approximately 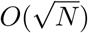 sparse, and furthermore that such matrices are also close to matrices with 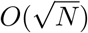 unique rows. These two tricks will allow us to compute sampling probabilities from population probabilities in *O*(*N*) time.

To show that the rows of **S** are approximately sparse we again use Hoeffding’s celebrated inequality. In his original paper, Hoeffding shows that his bounds on the tails of Binomial distributions also hold for Hypergeometric distributions (but phrased as sampling with and without replacement) [30]. We include the result below for completeness.

#### Lemma 9.

*(Adapted from [30, Theorem 4]) Let X* ∼ *H*_*N,K,n*_, *then for k* ≤ *nK/N*,

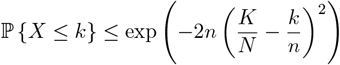

*and for k* ≥ *nK/N*,

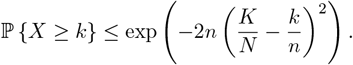

This lemma immediately implies an analogous sparsity result to Lemma 6. As the proof is essentially identical to the proof of that lemma we omit it.

#### Lemma 10.

*Let*

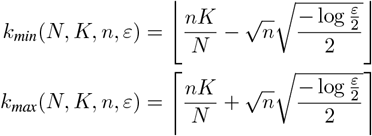

*and define* 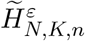 *as the distribution obtained by conditioning H*_*N,K,n*_ *to take values between k*_*min*_ + 1 *and k*_*max*_ − 1. *Then*,

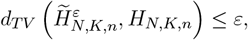

*and the mass function for* 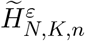 *contains only* 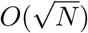 *non-zero entries*.

We will also need a result analogous to Lemma 2, showing that nearby Gypergeometric distributions are close in total variation distance. It turns out that the Hypergeometric case is slightly more delicate than the Binomial case. Our results rely on the assumption that we are not too close to sampling the entire population in that we will assume that *n ≤ αN* for some *α <* 1, and we will consider the *N* limit. Note that this still encompasses cases where we sample say 99% of the population, but rules out some pathological asymptotics such as having *n* = *N* − *N* ^1−*ε*^ where the proportion of the population sampled increases with the population size.

Before proving our general result about the total variation distance between different Hypergeometric distributions, we first prove a “one-step” inequality.

#### Lemma 11.

*Suppose that n ≤ αN for some α <* 1. *Furthermore, suppose that K ≤ N/*2. *Then, for any N larger than some finite N*_0_,

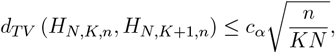

*where c*_*α*_ *is a universal constant that depends on α but is independent of n, N, and K*.

*Proof*. We start with the definition of total variation distance. Letting *p*_*k*_ be the probability that a random variable distributed as *H*_*N,K,n*_ is *k*, and letting *q*_*k*_ be the analogous quantity for *H*_*N,K*+1,*n*_ we have

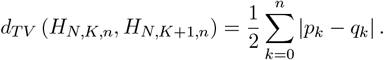

It is difficult to directly bound this sum sufficiently tightly, so instead we would like to convert it into an expectation by noting that

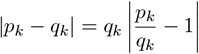

but an issue arises in that in general *H*_*N,K,n*_ and *H*_*N,K*+1,*n*_ can each put zero mass on an event where the other does not (i.e., neither is absolutely continuous with respect to the other), making the above potentially ill-defined. In particular, *q*_*k*_ is zero but *p*_*k*_ is still positive when *k* = *n* + *K N*.

Fortunately, we can use Lemma 9 to show that these events do not contribute substantially to the total variation distance. That is, if we take

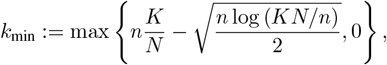

we can show that *k*_min_ *> n* + *K* − *N* by noting that

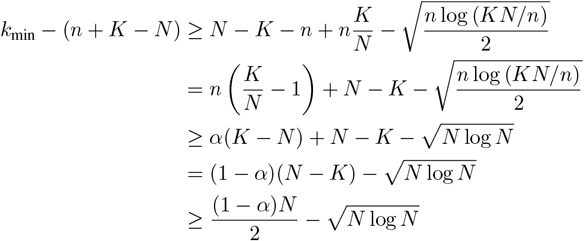

where we used that *n* ≤ *αN* and 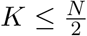 by assumption. The 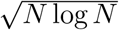 terms is lower order than (1− *α*)*N/*2, so 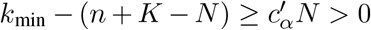 for some 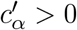 so long as *N* is large enough. This allows us to avoid the issue of dividing by zero, while only incurring an 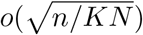 error term on the total variation, by Lemma 9:

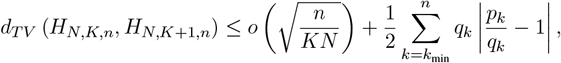

where we abuse notation and define the summand to be zero when *p*_*k*_ and *q*_*k*_ are both zero. Plugging in the values of *p*_*k*_ and *q*_*k*_ shows that when *q*_*k*_ is nonzero,

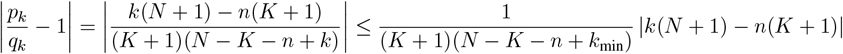

This allows us to bound the total variation distance in terms of an expectation of a random variable *X* distributed as *H*_*N,K*+1,*n*_.

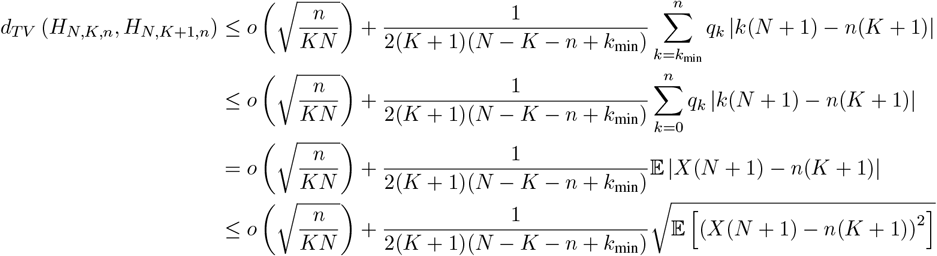

where the final line follows from Jensen’s inequality. This expectation only relies on the first two moments of a *H*_*N,K*+1,*n*_ random variable, and so may be readily, although tediously computed:

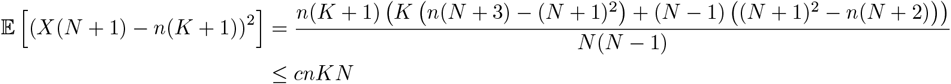

for some universal constant *c*, where we naively bounded all appearances of *n* and *K* beyond the first two terms in the numerator by factors of *N*. Noting from above that for *N* sufficiently large 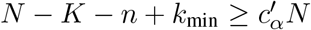 for some constant 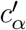 independent of *N, K*, and *n*, results in the desired bound:

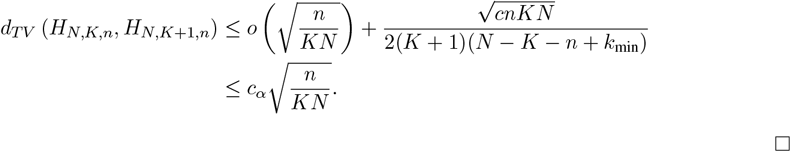

It then becomes quite easy to combine this one-step lemma to obtain something analogous to Lemma 3 but for Hypergeometric distributions.

#### Lemma 12.

*Suppose that n* ≤ *αN for some α <* 1 *and that* 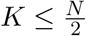. *There exists a constant c*_*α,ε*_ *that depends on α and ε but not N, K, or n such that for any non-negative integer* 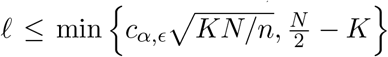 *we have*

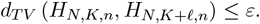

*Proof*. Total variation distance is a metric, and hence by the triangle inequality,

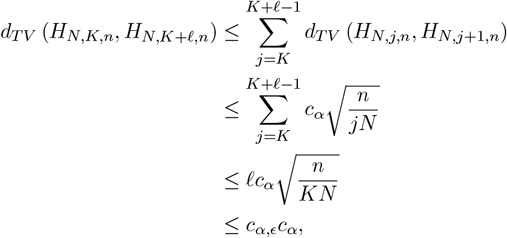

where the second line followed from Lemma 11. Choosing *c*_*α, ϵ*_ ≤ *ϵ/c*_*α*_ completes the proof.

We also need to consider where the first breakpoint can be.

#### Lemma 13.

*Suppose that n* ≤ *αN*. *Then, for any K such that K/N* ≤ (1 − *α*) 1 – [(1 − *ϵ*)^1*/n*^],

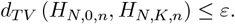

*In particular, for fixed ε, K/N may be as large as O*(1*/n*) *while maintaining a total variation distance of at most ε to the H*_*N*,0,*n*_ *distribution*.

*Proof*. If we let *X* ∼ *H*_*N,K,n*_, then, as in the proof of Lemma 1, the total variation distance between the distributions can be written as

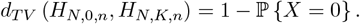

A quick calculation shows that

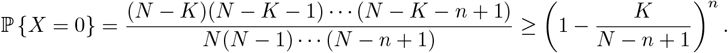

Using that *n* ≤ *αN*, we obtain

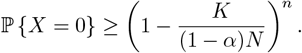

Therefore,

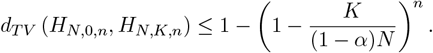

Solving for *K/N*, one obtains that the total variation distance is bounded by *ε* so long as

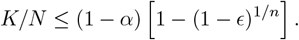

The term on the right hand side is the same, up to the factor of (1− *α*) as in the Binomial case. Therefore it is *O*(1*/n*) following the asymptotic argument in Lemma 1.

With Lemmas 12 and 13 in hand, we can prove a result analogous to Lemma 4 using similar techniques used in the proof of that result.

#### Lemma 14.

*Suppose that n* ≤ *αN for some α <* 1. *For fixed ε, there exist* 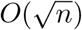 *breakpoints* 0 = *p*_0_ *< p*_1_ *< p*_2_ *< … < p*_*M*_ = 1 *such that for any K and K*′ *with K/N and K*′*/N being in the same block (i*.*e*., *there exists an i such that K/N, K*′*/N* ∈ [*p*_*i*_, *p*_*i*+1_]*) we have*

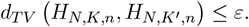

*Proof*. The proof follows immediately from the proof of Lemma 4, by noting that we may replace Lemma 3 by Lemma 12 and we may replace Lemma 1 by Lemma 13.

Combining these lemmas, we obtain our main approximation result for the sampling matrix.

#### Proposition 2.

*Let* **S** ∈ ℝ^*N* +1,*n*+1^ *be a Hypergeometric sampling matrix. We can build a representation of a matrix* 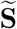 *with the following properties:*

- 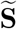 *has* 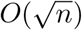 *unique rows*,
- *Each row of* 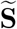 *has at most* 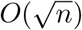 *non-zero elements*,
- 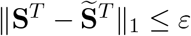.

*Furthermore, we can construct this representation in O*(*N*) *time and store it in O*(*N*) *space*.

*Proof*. The result follows immediately from an argument analogous to the proof of Proposition 1.

Finally, we note that this approximate matrix satisfies the properties of Lemma 8 allowing us to compute sample likelihoods from population likelihoods in *O*(*N*) time.

## B Practical considerations

In this section we will consider some practical aspects of using Binomial transition matrices in the context of the DTWF model. First, we will discuss a practical implementation detail that allows for faster likelihood computations when the underlying DTWF dynamics do not change too frequently. This trick will also allow us to quickly compute stationary distributions to a desired level of accuracy. We then discuss two aspects of the DTWF model that are useful in practice — an infinite sites version of the DTWF model to compute frequency spectra, and a version of the DTWF model conditioned on non-fixation, which is similar in spirit to the infinite sites model, but is conceptually cleaner when considering models with recurrent mutation.

### B.1 Faster repeated matrix-vector products

As discussed in the main text, we may need to repeatedly perform matrix vector products to compute likelihoods under the DTWF model for non-equilibrium populations. If the underlying dynamics of the DTWF model do not change too frequently, then we can obtain substantial computational savings by considering a “condensed” transition matrix and then repeatedly squaring that condensed matrix. In particular, we consider the case where we need to compute 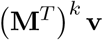 for large *k*. In principle, we could use Theorem 1 to approximate this by *k* matrix-vector products as 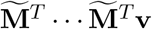 (evaluated right to left), which would cost *O*(*kN*) time, but if *k* is large we can speed this up substantially. One can view our fast multiplication algorithm for 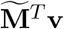, Equation 3, as consisting of two steps. First we project **v** into a 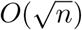 dimensional space by summing all of the entries of **v** that correspond to each of the 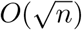 unique rows of 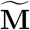, and then we multiply the resulting vector by the transpose of a matrix where we only keep non-redundant rows of 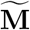, which we will call 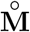. If we were to repeat this process, we would then project the vector 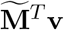 to the same 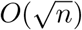 dimensional space. If we write this projection operation as a matrix, **Π**, then 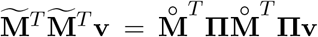. In general, 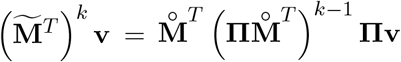. The trick is to note that 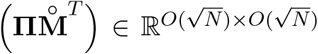 is a square matrix. As a result, we can repeatedly square 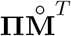, with each squaring taking *O*(*N* ^3*/*2^) time, allowing us to compute 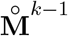 in *O*(*N* ^3*/*2^ log *k*) time. In principle this could be reduced substantially using faster matrix-matrix multiplication algorithms such as Strassen’s algorithm (reduces runtime to ≈ *O*(*N* ^1.4037^ log *k*)) [58] or the Coppersmith-Winograd algorithm (reduces runtime to ≈ *O*(*N* ^1.188^ log *k*)) [59]. Additionally, one could diagonalize the transition matrix and then compute matrix powers, which would require 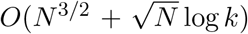 time. For simplicity and numerical stability we stick with the naive matrix-matrix multiplication algorithm, resulting in a runtime of *O*(*N* ^3*/*2^ log *k*). If *k* is *O*(*N*^*ε*+1*/*2^) or larger then this provides a faster algorithm. Similar tricks have been used in population genetics in the context of coalescent hidden Markov models [60, 61, 62].

### B.2 Computing the stationary distribution

Since we compute likelihoods forward in time, we must assume that at some point in the past the distribution of allele frequencies is known, and then use the DTWF transition matrix to integrate that distribution forward in time to the present. A natural choice is to assume that the population was at equilibrium at some point in the past, and to then compute the corresponding stationary distribution. By definition, when the system is at equilibrium, it is unchanged by the dynamics of the process, resulting in the following matrix equation:

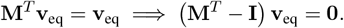

One could in principle solve this matrix equation (with the constraint that **v**_eq_ sum to one), but the naive strategy to obtain a solution costs *O*(*N* ^3^) time. We might hope that by simply replacing **M** by our approximation, 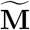 we might be able to solve this equation faster. While there are solvers that can take advantage of the sparsity of 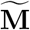, there are no solvers that can also take advantage of the fact 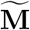 [63], only has a small number of unique rows. Here we propose two solutions, both of which rely on the ideas presented in Section B.1.

One solution is to solve for the equilibrium of the “condensed” dynamics. In the notation of Section B.1, we solve for 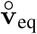 in

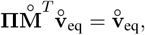

which requires *O*(*N*^3/2^) time. We then claim that 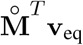 is an equilibrium of 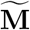. To see this, note that

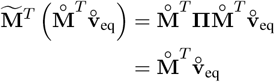

showing that 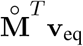 is invariant under the dynamics of 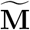.

This approach requires a particular choice of representative success probabilities when constructing 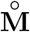. In principle, we would like to choose success probabilities using our moment matching approach, as described in the main text, but that requires knowing the equilibrium frequencies — exactly the object we wish to calculate. In practice, we find that using an iterative method where we use some initial guess of the equilibrium frequencies to construct 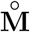, solve the above equation to obtain a new guess for the equilibrium frequencies, reconstruct 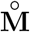, and so on, works well, resulting in errors on the order of machine precision after two or three iterations.

An alternative approach is to use the power method, which essentially approximates the equilibrium of a Markov chain by running the dynamics for a long time. That is,

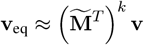

for any initial distribution, **v**, and taking *k* to be large. This is exactly the setting of Section B.1, and so we may use those results to speed up this computation. In practice, we can choose the initial **v** to be close to the true equilibrium by using analytical solutions to the Wright-Fisher Diffusion.

### B.3 Infinite sites

The mutation rate at any given position in the genome can be vanishingly small. For example, in humans the pre-modern effective population size was on the order of 10,000 [7], and per-base mutation rates are, on average, about 10^−8^ per generation [64]. Thus, one might expect to wait on the order of tens of thousands of generations (or hundreds of thousands of years) for a mutation to appear in the population at all. Furthermore, when a mutation first arrives in the population it arrives on only a single chromosome and as a result is likely to be quickly lost to drift. Under neutrality, and ignoring recurrent mutation, the DTWF model is a martingale, which means that the probability that a variant present is a single haploid ultimately fixes is one over the total number of haploids in the population. As a result, in order to get a mutation to fix at a site, it will on average require a mutation arising at that site a number of times equal to the population size. The waiting time between each of these mutations is on the order of hundreds of thousands of years. Ultimately, this means that at a single position, the timescale of equilibration is on the order of one over the mutation rate. In humans this would correspond to about a hundred million of generations, or a few billion years. Given that life has only been present on earth for about four billion years, and has clearly been rapidly changing, it seems implausible that any position in any genome could possibly be at equilibrium.

The infinite sites approximation avoids this issue of time-scales by approximating the genome as being infinitely long, with each site having an infinitesimally small mutation rate. In this limit, there are infinitely many sites, but only finitely many of them have variants segregating at any time, and as such it does not make sense to consider the probability of a given site segregating, as for any single site that probability is zero. Yet, one can model how many sites are expected to be segregating, or the distribution of allele frequencies conditioned on a site being segregating. Throughout, we will say the *frequency spectrum* to refer to the expected number of sites that have variants at each frequency. In this approximation, mutations only happen at most once at a given site and as a result we can determine which allele is “ancestral” and which allele is “derived”, and we can ignore recurrent mutations. One way to view the frequency spectrum is as a vector, *ϕ*, where *ϕ* (*k*) is the expected number of sites at which *k* individuals have the derived allele. We only track the dynamics of segregating sites (because there are always infinitely many non-segregating sites), and as such, we ignore any derived alleles if and when they reach fixation, so *ϕ* is *N*−1 dimensional.

Let *ϕ*_*t*_ denote the frequency spectrum at generation *t*. Alleles already in the population evolve according to DTWF dynamics, and we expect to “inject” *θ/*2 new singletons each generation, which we can capture by adding (*θ/*2)**e**_1_ to the frequency spectrum each generation, where **e**_1_ is the vector with 1 in the first position and zero in all other positions. One can then either add the mutations first, which would correspond to a model where mutation happens during gamete formation, and then the next generation is formed by sampling from those mutated gametes:

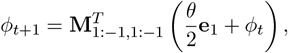

or one can consider a model where the next generation is formed first, and then mutates prior to being genotyped:

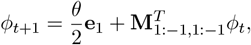

where the subscripts on **M** indicate that we are dropping the first and last rows and columns of **M** (corresponding to the non-segregating states). In the diffusion limit, generations happen infinitely fast so these two models are equivalent, but in the DTWF model these two mutation models are subtly different. In particular, mutating after demographic sampling produces substantially more singletons *in the population* — it is possible to show that the equilibrium of the model with mutation after demographic sampling will have exactly *θ/*2 more singletons in the population than the model with mutation before demographic sampling. Since the number of singletons in the equilibrium DTWF model where mutation happens after demographic sampling is ≈1.12 *×θ* [22], this results in a nearly factor of two difference between the models in terms of singletons. This difference is largely washed out in small subsamples of the population but becomes apparent as the sample size gets large. In humans there is some biological evidence for both of these models — siblings can share mutations that are not present in either parent, consistent with mutation in the parental germline, but there is also growing evidence that the first few replications after zygote formation are particularly error prone, which would be consistent with the second model [65, 66, 67]. Reality likely involves some combination of these models, indicating that the exact singleton count in large samples is not reliable, as it depends on extremely fine-scale aspects of the underlying model that are not currently well-understood. Yet, these details do not appear at all in the Wright-Fisher diffusion, providing further evidence that the diffusion stops providing a good approximation to reality as the sample size gets large.

Note that in either formulation, since we are dropping the first and last columns of **M**, we lose mass from *ϕ* each generation, corresponding exactly to those variants that have either reached fixation or been lost by drift. This loss of mass will at equilibrium be offset by the influx of mass from the (*θ/*2)**e**_1_ term corresponding to new mutations.

Since the above formulation is written entirely in terms of matrix-vector products, it is then easy to perform rapid approximate calculations using our tricks by simply replacing **M** with 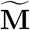.

Note that these formulations implicitly assume that the number of sites where a new mutation arises each generation is deterministic. A more realistic formulation would have a Poisson number of new mutations

– any given mutation is extremely unlikely, but there are many potential mutations throughout the genome, and so a “law of rare events” type argument gives the Poisson distribution. Yet, the variance of the Poisson equals the mean, so the average distance between an observation and the mean is about the square root of the mean. In our setting, that means that if the expected number of mutations per generation is large, then the Poissonian noise about that mean is comparatively small. For example, returning to the example of humans with an effective size of ≈10,000, a mutation rate of ≈10^−8^, and a genome size of ≈10^9^, we expect to see about 10,000 *×* 10^−8^ *×* 10^9^ = 10^5^ newly mutated sites per generation. Meanwhile, we expect the fluctuations in the number of mutations to be on the order of ≈1% of the expected number of mutations, showing that this deterministic approximation is quite accurate.

### B.4 Conditioning on non-fixation

The infinite sites model has some conceptual downsides. For example, methylated CpG sites have extremely high mutation rates [64, 25], making recurrent mutation an empirically non-negligible force at realistic population sizes [37]. Yet, the infinite sites model rules out recurrent mutation — if the mutation rate is high enough for a given site to mutate twice, then there must be infinitely many mutations in the genome. Furthermore, the notion of an “ancestral” and “derived” allele becomes unclear in the presence of recurrent mutation; if mutations can happen repeatedly at a given site then the derived allele could previously have gone to fixation and the ancestral allele could be subsequently reintroduced. A related conceptual issue is in the probability of a site being non-segregating. Under the infinite sites model, any given site is non-segregating with probability one, and the same holds for any finite number of sites. This prevents the infinite sites model from using the absence of mutations as an indication of natural selection, which has proven to be a powerful technique for measuring gene constraint [27, 25].

An idea that is conceptually similar to the infinite sites model is to allow arbitrary dynamics (e.g., recurrent mutation) but then condition the derived allele on non-fixation. That is, the derived allele is allowed to arise, perhaps even repeatedly at the same site, but we ignore situations in which that derived allele eventually fixes in the population. As a result, if we look at any position in the genome under this model, that position is either non-segregating with only the ancestral allele present, or it is segregating, but we know that at some point in the past it was non-segregating and the population was fixed for the ancestral allele. By making this assumption, we may pick a particular allele as being the ancestral allele, and safely know that under this model, the last time the population was monomorphic at this site, it was monomorphic with the ancestral allele. Yet, by allowing non-infinitesimal mutation rates at each position, we can still derive a probability of segregating for each site, and we can incorporate recurrent mutation in a conceptually clean way.

To incorporate conditioning on non-fixation, we can simply perform matrix-vector products as above, but we replace the final row of 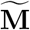(corresponding to the derived allele being fixed) with a row of zeros, and we replace the final column of 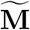 (corresponding to transitioning to having the derived allele being fixed) with a column of zeros. This causes any mass that would have resulted in fixation being removed from the system. As a result, each transition, 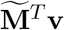, will result in a loss of mass corresponding to the allele trajectories that would have resulted in fixation of the derived allele. After each transition we can then renormalize **v** to sum to one. That this is correct follows from noting that we are eliminating all trajectories that result in fixation, and then rescaling all of the remaining probabilities by a constant (the probability of not reaching fixation).

## C Convergence of moments

In this section we consider the moments of the DTWF process as well as our approximation. While we showed in Appendix A that the processes are close in terms of total variation distance, total variation distance can be either overly strict or overly lax in some situations. This arises because total variation is agnostic to any metric on the space of outcomes. Our approximation produces a small difference in total variation distance, but there are other approximations that also have small total variation distance, but result in pathological behavior. For example, consider the process obtained by flipping a coin that comes up heads with probability *ε*, and if the coin is heads, then in the next generation the frequency of the *A* allele is zero regardless of its current frequency. If the coin is tails, then the next generation is obtained via the standard DTWF process. It is easy to see that the transition density of this strange process has small total variation distance to the transition density of the usual DTWF process. Yet, this construction has totally outlandish behavior — if we consider a DTWF model without recurrent mutation, and say that the current allele frequency is 1, then under the true DTWF model, the population will be forever stuck at a frequency of 1. On the other hand, the strange construction will eventually (after about 1*/ε* generations on average) crash to a frequency of 0. This example highlights that being close in total variation is not necessarily sufficient for one process to be a sensible approximation of the other. As such, the remainder of this section will work toward showing that the mean and variance of the transition density of our approximate process are close to the transition density of the full DTWF process.

There are two pieces to our approximation — combining rows of the transition matrix that correspond to Binomial distributions with similar success probabilities, and sparsifiying the row — and both affect the moments.

The effect of combining similar rows is straightforward to analyze. Since the mean of a *B*_*N,p*_ distribution is *Np*, and, assuming that *p* ≤ 1*/*2, we combine rows a row with success probability *p* with another that differs by at most 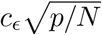, we can see that this affects the mean by 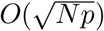. In practice, we also choose our representative success probabilities so that while some rows have their means increased by as much as 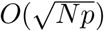, those are balanced by rows that have their means decreased by a corresponding amount. As a result, if you pick a row randomly (with probability proportional to the probability of observing that frequency in the previous generation) the difference between its approximate and true means is 0 on average.

Likewise, the variance of *B*_*N,p*_ is *Np*(1−*p*), which is also altered by something that is 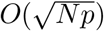.In both cases, we see that the effect of the perturbation is a lower order term.

Analyzing how the sparsification affects the moments is substantially more technical, but we include it here for completeness. We will prove the closeness of the first two moments of the truncated and non-truncated Binomial distributions by showing that the truncated Binomial distribution is very close to a truncated Normal distribution, for which we can readily compute moments, and then show that those moments are close to those of the Binomial distribution. Our proof relies on a non-uniform version of the celebrated Berry-Esseen theorem [68], which we include here without proof.

### Lemma 15.

*(Non-uniform Berry-Esseen bound for Binomial, adapted from [68, Theorem 3]) Let X ∼ B*_*N,p*_, *and let Z ∼ 𝒩* (0, 1). *Then, there exists a universal constant c such that*

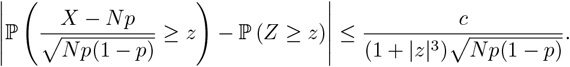

Using the Berry-Esseen bound we can show that Binomial distributions truncated as described in Lemma 6 (and used in our approach) are quantitatively similar in distribution to a truncated Gaussian.

### Lemma 16.

*(Non-uniform Berry-Esseen bound for truncated Binomial) Let* 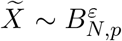 *be a random variable*

*drawn from a Binomial distribution truncated as described in Lemma 6. Define* 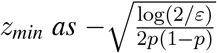, *and* 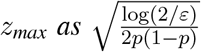. *Let* 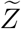 *be distributed according to a truncated standard Normal distribution, truncated at z*_*min*_ *and z*_*max*_. *Then*,

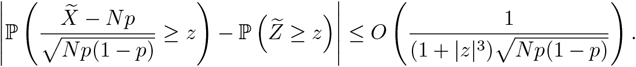

*Proof*. First, note that by construction 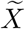 and 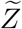 are truncated at the same points (that is *z*_min_ and *z*_max_ are simply the centered and scaled versions of *k*_min_ and *k*_max_ appearing in Lemma 6) so for any *z* lying outside of these truncations points, the bound in the Lemma is vacuously true, as the difference in the distributions is 0.

Now, consider a *z ∈* [*z*_min_, *z*_max_]. We begin by rewriting the distribution of 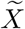 in terms of the distribution of a Binomial random variable, *X* ∼ *B*_*N,p*_:

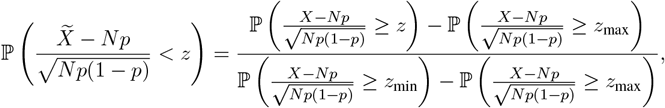

and using Lemma 15 we can write the right hand side as

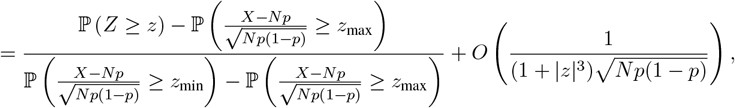

where we used that by Lemma 6 the denominator is at least some constant independent of *N, p*, and *z*. We now write the probability involving *Z* in terms of 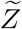

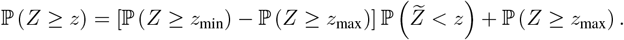

Plugging this result in, we obtain

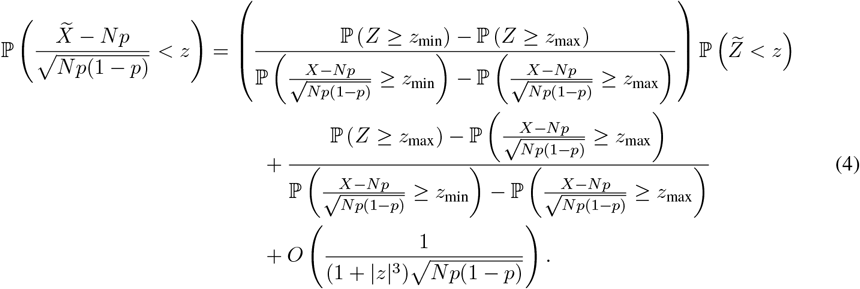

We now tackle the two terms on the right hand side of the previous equation, starting with the second term.

Bounding the denominator by a constant, as above, and applying Lemma 15 we see

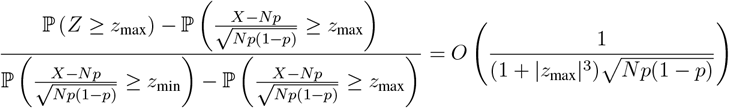

Now, noticing that since *z*∈[*z*_min_, *z*_max_], we have that |*z*| ≤|*z*_max_| by construction, and so we can loosen this bound to

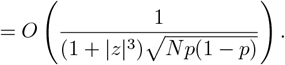

We now turn to the first term on the right hand side of Equation 4, where we use similar tricks:

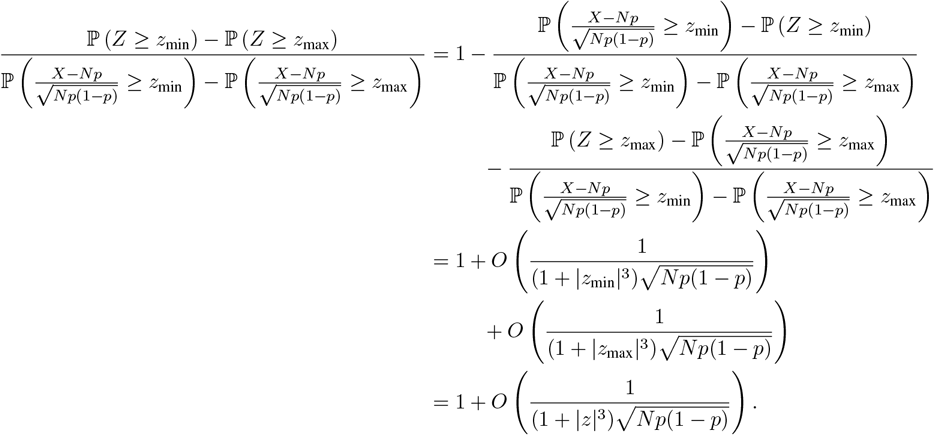

Using these results, we may simplify Equation 4 to obtain

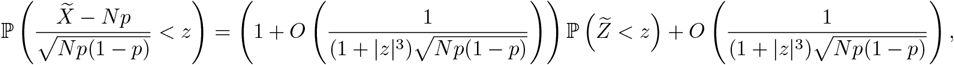

and noticing that 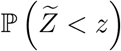 is bounded by 1 we finally obtain

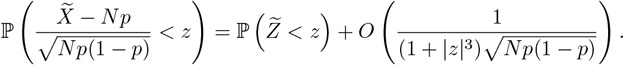

With our truncated version of the non-uniform Berry-Esseen theorem, we are now ready to prove that our truncation of the Binomial distribution does not substantially alter the first two moments.

### Lemma 17.

*Let* 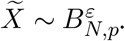 *Then*,

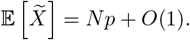

*Proof*. Recall that for any strictly positive random variable,

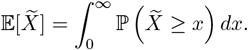

Centering and scaling, we see

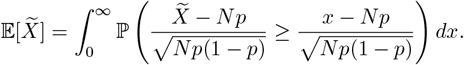

Performing a change of variables and applying Lemma 16, we arrive at

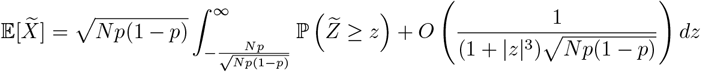

where 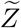 is a truncated Gaussian, truncated at 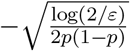 and 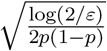.

Noting that

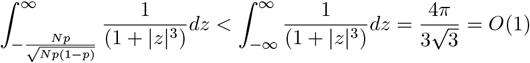

we obtain

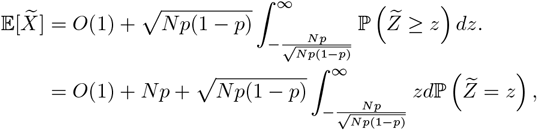

where the second equality follows from integrating by parts. Now, note that without loss of generality we can assume that *p* ≥ 1*/*2 (by symmetry of the Binomial). Therefore, if we take *N* to be large enough, the lower limit of the integral is lower than the lower truncation point of 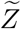 so the integral covers the entire domain of 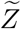. Therefore, for sufficiently large *N*, the integral is exactly 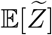. The mean of a truncated standard Gaussian with symmetric truncation points is 0, completing the proof.

### Lemma 18.

*Let* 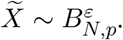. *Then*,

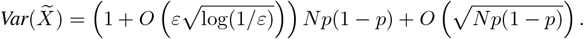

*Proof*. Let *μ*_*B*_ be the mean of 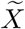. Then, by Lemma 17,

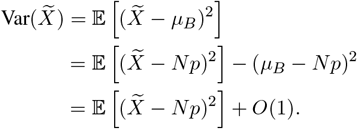

We now evaluate the expectation on the right hand side

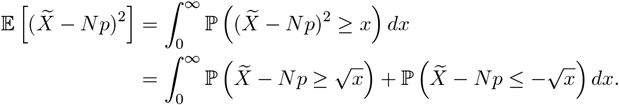

We now perform the change of variables 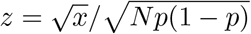, and use Lemma 16, with 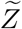 being the truncated Gaussian corresponding to 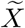 :

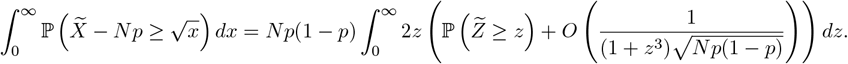

Noting that

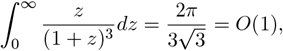

we obtain

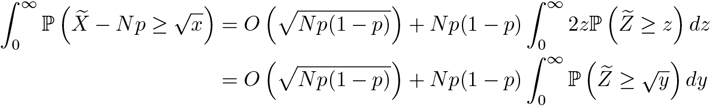

where the final line follows from the change of variables 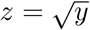. A similar computation shows that

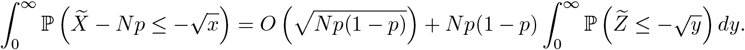

We therefore have that

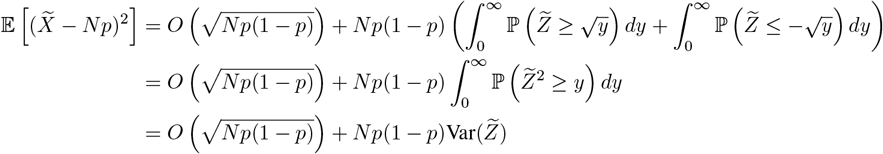

where the final line follows by noting that the melan of a symmetrically truncated standard Normal is 0. 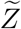 is a standard Normal symmetrically truncated at 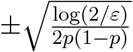. Letting *z*_min_ and *z*_max_ be these truncation points *ϕ* (·) and Φ(·) be the probability density function and cumulative density function of the standard Normal respectively, we have that

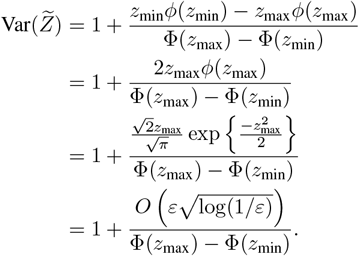

The denominator can be bounded from below by 1 − *O*(*ε*), completing the proof.

Together, these results show that in addition to being close in total variation distance, our approximate process is also close in terms of the first two moments.

## D Proof of the representation of the 1-operator norm

For completeness we include a proof that the 1-operator norm of a matrix is the max of the 𝓁_1_ norm across columns. This is a standard, well-known result.

*Proof*. Consider an *N*×*P* matrix *A*. We will complete the proof in two steps – first we will show that ‖*A*‖_1_ is at least as large as the max of the 𝓁_1_ norm across columns, then we will show that ‖*A*‖_1_ is no larger than the max of the 𝓁_1_ norm across columns.

Without loss of generality, assume that the first column of *A* has the largest 𝓁_1_ norm. Consider the vector

*e*_1_, which has a 1 for its first entry and zero for all other entries. Clearly ‖*e*_1_‖_1_ = 1, and so

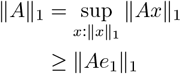

but ‖*Ae*_1_‖_1_ is just the 𝓁_1_ norm of the first column of *A* which is the column with the largest 𝓁_1_ norm.

To prove the other direction, we will need to consider the columns of *A* which we will write as *A*_·,1_, …, *A*_·,*P*_. We see that for any *x* with ‖*x*‖_1_ = 1,

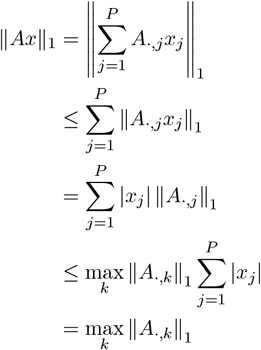

where the first inequality followed from the triangle inequality.

## E Additional numerical results

Here we include additional details, plots, and results related to Section 3.2. Throughout Section 3.2 we use slightly modified versions of the demographic histories for the CEU and YRI samples from the 1000 Genomes Project [36] inferred using MSMC [7]. First, we smoothed out some small fluctuations in sizes older than one hundred thousand years ago, and assumed that an extremely large population size estimated using YRI but not estimated using CEU was artifactual, opting to use a smaller ancestral size. Second, the present day YRI population size (36,822) is smaller than the largest sample sizes we wanted to consider, so we added a single generation of 500,000 individuals at present. Years were converted to generation times by dividing by 30 and rounding. The original, and modified demographies (ignoring the recent increase in YRI) are presented in Figure A1.

Modifying the YRI demography this way results in an extreme scenario where many individuals must find a common ancestor one generation ago since a sample size of *n* = 300,000 is ≈10× larger than the population size in the previous generation. An interesting consequence of this is that under neutrality there can be many *de novo* mutations occurring in the most recent generation resulting in singletons, but since the expected number of present day haplotypes that come from a particular parent in the previous generation is ≈10, variants that are present in more than one but fewer than 10 individuals should be exceedingly rare. As a result, the frequency spectrum under this demography is non-convex: doubletons are rarer than both singletons and tripletons, but more common than extremely high frequency variants (Figure A2). This highlights an advantage of our approach in that we can model such unusual frequency spectra whereas the frequency spectrum under the diffusion approximation must be convex for *any* population size history, a result of Sargsyan and Wakeley [69]. We note, however, that this particular result is driven by our simplistic modification of the YRI demography and unlikely to be a good match to actual frequency spectra obtained from large samples of individuals closely related to the YRI sample from the 1000 Genomes Project. Yet, for small sample sizes, we see that the frequency spectra are largely unaffected by this single generation of growth, and the frequency spectra take on more familiar shapes.

**Figure A1:**
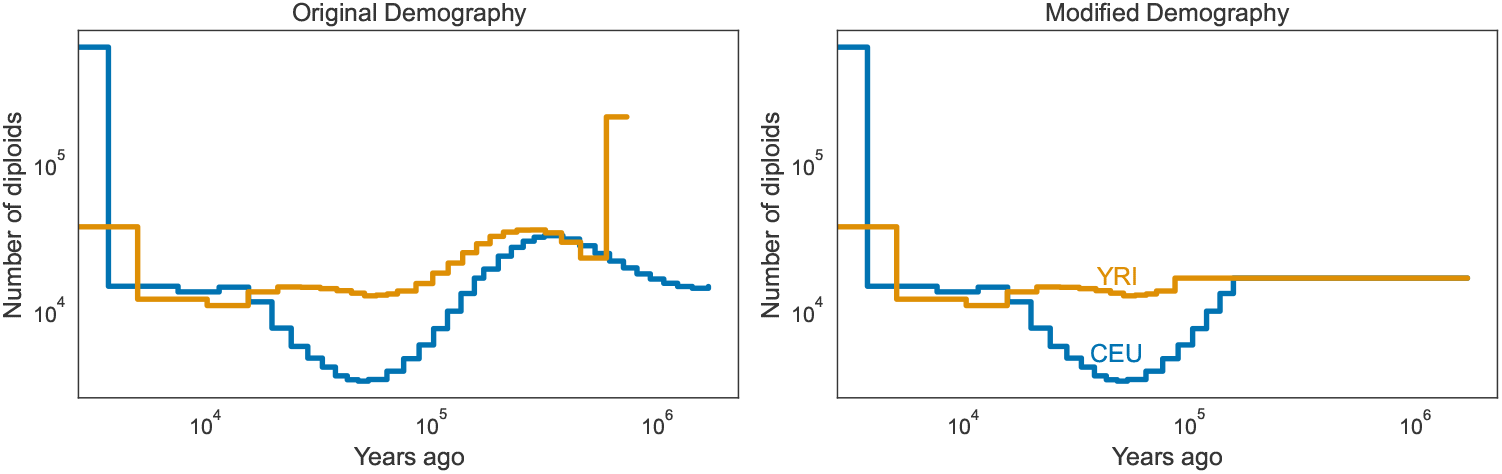
The original [7] and modified demographies for CEU and YRI considered in this paper. Note that in the modified demography, YRI has an additional generation with a population size of 500,000 diploids in the most recent generation that is truncated from the plot.

To compute the Fisher Information in Section 3.2 we used numerical differentiation on linear interpolations of likelihoods. Specifically, we compute likelihoods by linearly interpolating between precomputed grid points. This method results in slight technical artifacts where the Fisher Information is extremely high on one side of a grid point and extremely low on the other side of the grid point with a discontinuity at the grid point. This arises from linear interpolation being non-differentiable at the grid points. To avoid these technical artifacts, we computed the Fisher Information on a dense grid of points and then smoothed the resulting values using `gaussian filter1d` from `scipy` [63] with a kernel chosen to visually smooth out the artifactual fluctuations in the Fisher Information.

**Figure A2:**
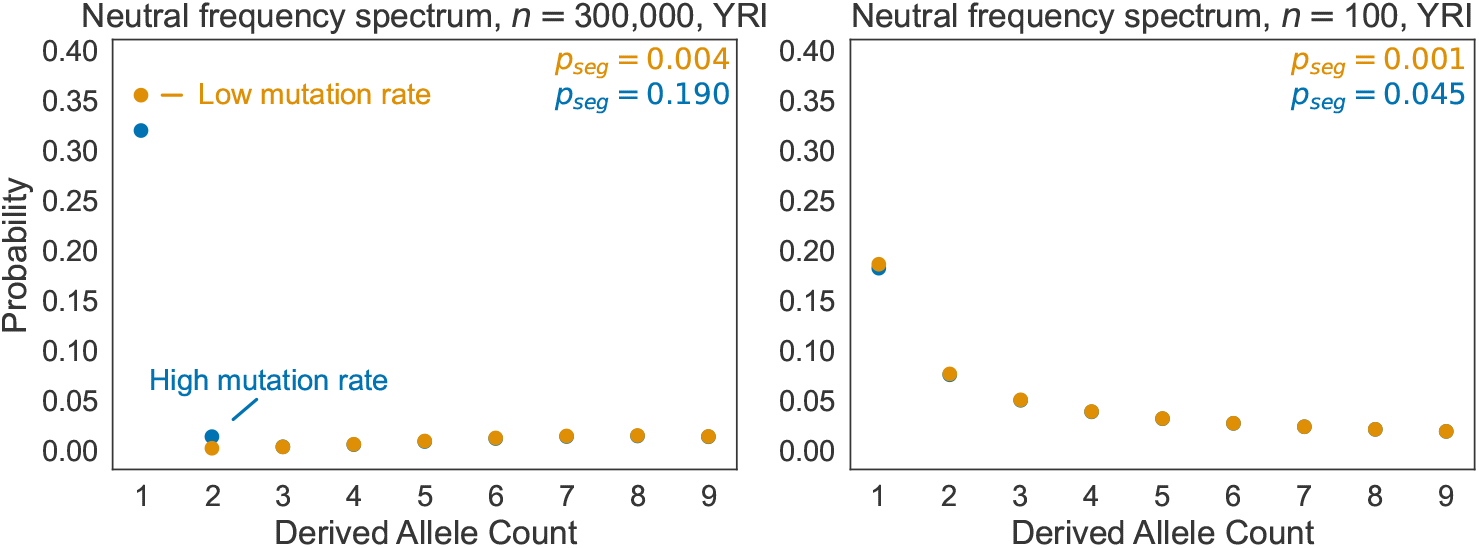
The lowest entries of the frequency spectra implied by our modified YRI demography for sample sizes of *n* = 300,000 diploids (left) or *n* = 100 diploids (right). The high mutation rate corresponds to the mutation rate of a methylated CpG (1.25×10^−7^ per generation) and the low mutation rate roughly corresponds to the rate of transversions (2.44×10^−9^ per generation). The spectra for the two mutation rates almost coincide on the right plot.

## Notes

### Competing Interest Statement

The authors have declared no competing interest.

## References

[1] Gillespie JH. Population genetics: a concise guide. JHU press; 2004.

[2] Ewens WJ. Mathematical population genetics: theoretical introduction. vol. 27. Springer; 2004.

[3] Sawyer SA, Hartl DL. Population genetics of polymorphism and divergence. Genetics. 1992;132(4):1161–1176.

[4] Bhaskar A, Wang YR, Song YS. Efficient inference of population size histories and locus-specific mutation rates from large-sample genomic variation data. Genome Research. 2015;25(2):268–279.

[5] Gutenkunst RN, Hernandez RD, Williamson SH, Bustamante CD. Inferring the joint demographic history of multiple populations from multidimensional SNP frequency data. PLoS Genetics. 2009;5(10):e1000695.

[6] Kim BY, Huber CD, Lohmueller KE. Inference of the distribution of selection coefficients for new nonsynonymous mutations using large samples. Genetics. 2017;206(1):345–361.

[7] Schiffels S, Durbin R. Inferring human population size and separation history from multiple genome sequences. Nature Genetics. 2014;46(8):919–925.

[8] Song YS, Steinrücken M. A simple method for finding explicit analytic transition densities of diffusion processes with general diploid selection. Genetics. 2012;190(3):1117–1129.

[9] Steinrücken M, Wang YR, Song YS. An explicit transition density expansion for a multi-allelic Wright– Fisher diffusion with general diploid selection. Theoretical Population Biology. 2013;83:1–14.

[10] Živković D, Steinrücken M, Song YS, Stephan W. Transition densities and sample frequency spectra of diffusion processes with selection and variable population size. Genetics. 2015;200(2):601–617.

[11] Kamm JA, Terhorst J, Song YS. Efficient computation of the joint sample frequency spectra for multiple populations. Journal of Computational and Graphical Statistics. 2017;26(1):182–194.

[12] Kingman JFC. The coalescent. Stochastic Processes and their Applications. 1982;13(3):235–248.

[13] Jansen S, Kurt N. On the notion(s) of duality for Markov processes. Probability Surveys. 2014;11(none):59–120. Available from: https://doi.org/10.1214/12-PS206.

[14] Fu YX. Exact coalescent for the Wright–Fisher model. Theoretical Population Biology. 2006;69(4):385– 394.

[15] Polanski A, Kimmel M. New explicit expressions for relative frequencies of single-nucleotide polymorphisms with application to statistical inference on population growth. Genetics. 2003;165(1):427–436.

[16] Krone SM, Neuhauser C. Ancestral processes with selection. Theoretical Population Biology. 1997;51(3):210–237.

[17] Kamm J, Terhorst J, Durbin R, Song YS. Efficiently inferring the demographic history of many populations with allele count data. Journal of the American Statistical Association. 2020;115(531):1472–1487.

[18] Jouganous J, Long W, Ragsdale AP, Gravel S. Inferring the joint demographic history of multiple populations: beyond the diffusion approximation. Genetics. 2017;206(3):1549–1567.

[19] Bhaskar A, Clark AG, Song YS. Distortion of genealogical properties when the sample is very large. Proceedings of the National Academy of Sciences. 2014;111(6):2385–2390.

[20] Krukov I, Gravel S. Taming strong selection with large sample sizes. bioRxiv. 2021; p. 2021–03.

[21] Melfi A, Viswanath D. Single and simultaneous binary mergers in Wright-Fisher genealogies. Theoretical Population Biology. 2018;121:60–71.

[22] Wakeley J, Takahashi T. Gene genealogies when the sample size exceeds the effective size of the population. Molecular Biology and Evolution. 2003;20(2):208–213.

[23] Agarwal I, Fuller ZL, Myers SR, Przeworski M. Relating pathogenic loss-of function mutations in humans to their evolutionary fitness costs. eLife. 2023;12:e83172.

[24] Cassa CA, Weghorn D, Balick DJ, Jordan DM, Nusinow D, Samocha KE, et al. Estimating the selective effects of heterozygous protein-truncating variants from human exome data. Nature Genetics. 2017;49(5):806–810.

[25] Karczewski KJ, Francioli LC, Tiao G, Cummings BB, Alfoöldi J, Wang Q, et al. The mutational constraint spectrum quantified from variation in 141,456 humans. Nature. 2020;581(7809):434–443.

[26] LaPolice TM, Huang YF. A deep learning framework for predicting human essential genes from population and functional genomic data. bioRxiv. 2021; p. 2021–12.

[27] Lek M, Karczewski KJ, Minikel EV, Samocha KE, Banks E, Fennell T, et al. Analysis of protein-coding genetic variation in 60,706 humans. Nature. 2016;536(7616):285–291.

[28] Weghorn D, Balick DJ, Cassa C, Kosmicki JA, Daly MJ, Beier DR, et al. Applicability of the Mutation– Selection Balance Model to Population Genetics of Heterozygous Protein-Truncating Variants in Humans. Molecular Biology and Evolution. 2019;36(8):1701–1710.

[29] Zeng T, Spence JP, Mostafavi H, Pritchard JK. Bayesian estimation of gene constraint from an evolutionary model with gene features. bioRxiv. 2023;.

[30] Hoeffding W. Probability Inequalities for Sums of Bounded Random Variables. Journal of the American Statistical Association. 1963;58(301):13–30. Available from: https://www.tandfonline.com/doi/abs/10.1080/01621459.1963.10500830.

[31] Krukov I, de Sanctis B, de Koning APJ. Wright–Fisher exact solver (WFES): scalable analysis of population genetic models without simulation or diffusion theory. Bioinformatics. 2016 12;33(9):1416–1417. Available from: https://doi.org/10.1093/bioinformatics/btw802.

[32] Tataru P, Simonsen M, Bataillon T, Hobolth A. Statistical Inference in the Wright–Fisher Model Using Allele Frequency Data. Systematic Biology. 2016 08;66(1):e30–e46. Available from: https://doi.org/10.1093/sysbio/syw056.

[33] Spence JP, Song YS. Inference and analysis of population-specific fine-scale recombination maps across 26 diverse human populations. Science Advances. 2019;5(10):eaaw9206.

[34] Bustamante CD, Wakeley J, Sawyer S, Hartl DL. Directional selection and the site-frequency spectrum. Genetics. 2001;159(4):1779–1788.

[35] Melfi A, Viswanath D. The Wright–Fisher site frequency spectrum as a perturbation of the coalescent’s. Theoretical Population Biology. 2018;124:81–92.

[36] Consortium GP, et al. An integrated map of genetic variation from 1,092 human genomes. Nature. 2012;491(7422):56.

[37] Harpak A, Bhaskar A, Pritchard JK. Mutation rate variation is a primary determinant of the distribution of allele frequencies in humans. PLoS Genetics. 2016;12(12):e1006489.

[38] Wakeley J, Fan WTL, Koch E, Sunyaev S. Recurrent mutation in the ancestry of a rare variant. bioRxiv. 2022;Available from: https://www.biorxiv.org/content/early/2022/08/18/2022.08.18.504427.

[39] Agarwal I, Przeworski M. Mutation saturation for fitness effects at human CpG sites. eLife. 2021;10:e71513.

[40] Backman JD, Li AH, Marcketta A, Sun D, Mbatchou J, Kessler MD, et al. Exome sequencing and analysis of 454,787 UK Biobank participants. Nature. 2021;599(7886):628–634.

[41] Adrion JR, Cole CB, Dukler N, Galloway JG, Gladstein AL, Gower G, et al. A community-maintained standard library of population genetic models. eLife. 2020;9:e54967.

[42] Kamm JA, Spence JP, Chan J, Song YS. Two-locus likelihoods under variable population size and finescale recombination rate estimation. Genetics. 2016;203(3):1381–1399.

[43] Diaz-Papkovich A, Anderson-Trocmé L, Ben-Eghan C, Gravel S. UMAP reveals cryptic population structure and phenotype heterogeneity in large genomic cohorts. PLoS Genetics. 2019;15(11):e1008432.

[44] Yengo L, Robinson MR, Keller MC, Kemper KE, Yang Y, Trzaskowski M, et al. Imprint of assortative mating on the human genome. Nature Human Behaviour. 2018;2(12):948–954.

[45] Cannings C. The latent roots of certain Markov chains arising in genetics: a new approach, I. Haploid models. Advances in Applied Probability. 1974;6(2):260–290.

[46] Pitman J. Coalescents with multiple collisions. Annals of Probability. 1999; p. 1870–1902.

[47] Eldon B, Wakeley J. Coalescent processes when the distribution of offspring number among individuals is highly skewed. Genetics. 2006;172(4):2621–2633.

[48] Mohle M, Sagitov S. A classification of coalescent processes for haploid exchangeable population models. Annals of Probability. 2001; p. 1547–1562.

[49] Spence JP, Kamm JA, Song YS. The site frequency spectrum for general coalescents. Genetics. 2016;202(4):1549–1561.

[50] Jewett EM, Steinrücken M, Song YS. The effects of population size histories on estimates of selection coefficients from time-series genetic data. Molecular Biology and Evolution. 2016;33(11):3002–3027.

[51] Steinrücken M, Bhaskar A, Song YS. A novel spectral method for inferring general diploid selection from time series genetic data. The Annals of Applied Statistics. 2014;8(4):2203.

[52] Mathieson I, Terhorst J. Direct detection of natural selection in Bronze Age Britain. Genome Research. 2022;32(11-12):2057–2067.

[53] Paszke A, Gross S, Massa F, Lerer A, Bradbury J, Chanan G, et al. Pytorch: An imperative style, highperformance deep learning library. Advances in Neural Information Processing Systems. 2019;32.

[54] Gao Y, Liu Y, Zhang H, Li Z, Zhu Y, Lin H, et al. Estimating GPU memory consumption of deep learning models. In: Proceedings of the 28th ACM Joint Meeting on European Software Engineering Conference and Symposium on the Foundations of Software Engineering; 2020. p. 1342–1352.

[55] Chen RT, Rubanova Y, Bettencourt J, Duvenaud DK. Neural ordinary differential equations. Advances in Neural Information Processing Systems. 2018;31.

[56] Adell JA, Jodrá P. Exact Kolmogorov and total variation distances between some familiar discrete distributionsolmogorov and total variation distances between some familiar discrete distributions. Journal of Inequalities and Applications. 2006;2006:1–8.

[57] Roos B. Binomial approximation to the Poisson binomial distribution: The Krawtchouk expansion. Theory of Probability & Its Applications. 2001;45(2):258–272.

[58] Strassen V. Gaussian elimination is not optimal. Numerische Mathematik. 1969;13(4):354–356.

[59] Coppersmith D, Winograd S. Matrix multiplication via arithmetic progressions. In: Proceedings of the Nineteenth Annual ACM Symposium on Theory of Computing; 1987. p. 1–6.

[60] Paul JS, Song YS. Blockwise HMM computation for large-scale population genomic inference. Bioinformatics. 2012;28(15):2008–2015.

[61] Steinrücken M, Kamm J, Spence JP, Song YS. Inference of complex population histories using wholegenome sequences from multiple populations. Proceedings of the National Academy of Sciences. 2019;116(34):17115–17120.

[62] Terhorst J, Kamm JA, Song YS. Robust and scalable inference of population history from hundreds of unphased whole genomes. Nature Genetics. 2017;49(2):303–309.

[63] Virtanen P, Gommers R, Oliphant TE, Haberland M, Reddy T, Cournapeau D, et al. SciPy 1.0: fundamental algorithms for scientific computing in Python. Nature Methods. 2020;17(3):261–272.

[64] Jónsson H, Sulem P, Kehr B, Kristmundsdottir S, Zink F, Hjartarson E, et al. Parental influence on human germline de novo mutations in 1,548 trios from Iceland. Nature. 2017;549(7673):519–522.

[65] Gao Z, Moorjani P, Sasani TA, Pedersen BS, Quinlan AR, Jorde LB, et al. Overlooked roles of DNA damage and maternal age in generating human germline mutations. Proceedings of the National Academy of Sciences. 2019;116(19):9491–9500.

[66] Gao Z, Wyman MJ, Sella G, Przeworski M. Interpreting the dependence of mutation rates on age and time. PLoS Biology. 2016;14(1):e1002355.

[67] Sasani TA, Pedersen BS, Gao Z, Baird L, Przeworski M, Jorde LB, et al. Large, three-generation human families reveal post-zygotic mosaicism and variability in germline mutation accumulation. eLife. 2019;8:e46922.

[68] Nagaev SV. Some limit theorems for large deviations. Theory of Probability & Its Applications. 1965;10(2):214–235.

[69] Sargsyan O, Wakeley J. A coalescent process with simultaneous multiple mergers for approximating the gene genealogies of many marine organisms. Theoretical Population Biology. 2008;74(1):104–114.

